# Extracellular vesicle–associated nucleic acids and proteins as a fraction for fish biodiversity monitoring and physiological-signal recovery

**DOI:** 10.64898/2026.07.15.735187

**Authors:** Jafar Hayatov, Xueli Xu, Lu Long, Yangyu Wu, Ricardo David Avellan Llaguno, Alex Obeten Ujong, Zhizhen Pan, Qiansheng Huang

## Abstract

Monitoring the biodiversity and physiological condition of aquatic organisms under environmental stress is central to aquatic ecosystem management. Environmental nucleic acid methods (eDNA and eRNA) have transformed biomonitoring but face well-documented limitations, including susceptibility to degradation, difficulty distinguishing living from legacy or transported signals, and limited capacity to report organism condition. Extracellular vesicles (EVs) released from all kinds of species, are stable and widespread vesicles in environment, showing biomarker potential. Here we evaluate whether nucleic acids and proteins recovered from EVs provide a complementary capture fraction that helps address these limitations. All four targets (eDNA, eRNA, EV-DNA, EV-RNA) shared an identical 12S rRNA metabarcoding workflow and differed only in the pre-analytical capture step, filtration for bulk environmental nucleic acids versus sequential 0.22 µm filtration, tangential flow filtration, and ultracentrifugation for the EV-encapsulated fraction. In a controlled aquarium, EV-DNA recovered all seven reference species (100% sensitivity) versus 85.7% for bulk methods, including a low-abundance detection of *Anguilla japonica* that warrants independent confirmation. In field sampling at Xinglinwan Reservoir, EV-RNA recovered 11 of 12 expected species (91.7% sensitivity) compared with 50–58% for bulk methods, including four reference taxa detected only by the EV fraction. EV-based methods also captured more even community representation, and EV-RNA showed lower human-read contamination than eDNA. Metaproteomic analysis of reservoir EVs recovered fish-derived proteins, dominated by *Cyprinus carpio*, including candidate stress-associated functions (chaperones, metallothioneins, oxidoreductases), independently corroborating the EV-based detection of *C. carpio*. Their abundance varied with seasons, suggesting their ability to respond to environmental changes. Together, these results indicate that EV-associated nucleic acids and proteins constitute an informative complementary fraction for integrated biodiversity and physiological-signal monitoring in impacted aquatic environments.

**Graphical abstract:** 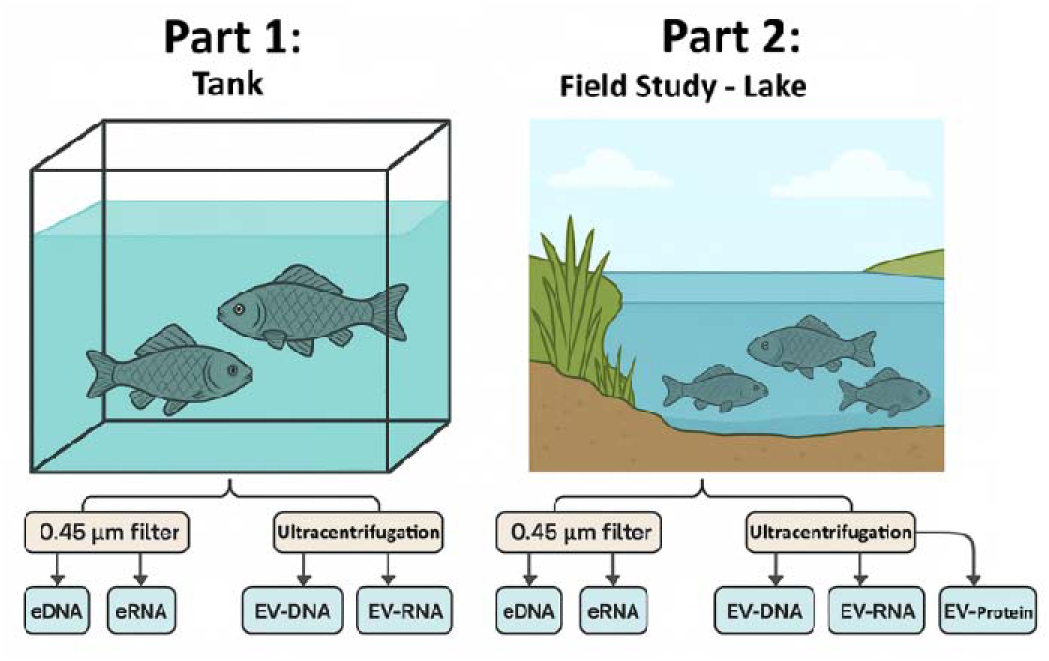

## 1. INTRODUCTION

Biomonitoring is the systematic observation and measurement of organisms or communities to assess ecosystem health, detect environmental changes, and support the management and protection of aquatic environments threatened by human activities^1^.

Environmental DNA and RNA (eDNA/eRNA) are free nucleic acids that exist independently in nature, originating from urine, feces, mucus, or shed cells, and can be collected from various environments such as water, soil, and air ^2,3^. In recent years, the use of eDNA and the growing popularity of eRNA have transformed this field. These two techniques have the potential to increase the temporal and spatial acuity of biomonitoring programs, with the latter approach having the capacity to significantly enhance our functional understanding of real-time organismal and environmental health^4^. In contrast to these, the only alternative method is traditional field survey methods, such as electrofishing, gill nets, and traps, followed by morphological identification, which are expensive, poorly reproducible, invasive, and require expertise in taxonomy^5^.

For eDNA and eRNA, we can observe that each has its strengths and weaknesses in various aspects. Unlike eDNA, eRNA can distinguish living organisms and assess population health^6^, but has limitations such as high sensitivity to pH and temperature changes^7^. DNA molecules have a more stable structure, making them inherently more resistant to such stress factors. In fact, under certain conditions, the stability of DNA molecules allows them to be preserved for millions of years, making them useful in ancient DNA studies ^8^. However, the long lifespan of DNA often creates challenges when it comes to false positives. For instance, in analyses conducted in rivers or dynamic water bodies, a DNA molecule from a distant location might be detected, or be carried by other living organisms, providing misleading information^9^. In contrast, because RNA molecules are short-lived, they do not provide these misleading information and instead offer data specific to the biodiversity of the area ^10^. However, the short lifespan of eRNA and its rapid degradation can lead to false-negative results. In certain cases, some researchers even try to combine these two methods at the metabarcoding stage to achieve more accurate results, in order to overcome the shortcomings of these two methods and ensure more effective detectability ^11^. Furthermore, while eRNA-based health assessment has shown promise in controlled laboratory exposures, no field-scale implementation in lakes or open water bodies has yet been demonstrated, leaving health monitoring as an unmet need in environmental biomonitoring practice.^10,12–14^

Cells produce extracellular vesicles (EVs) as a fundamental mechanism for intercellular communication, conserved throughout all three domains of life^15,16^. EVs-associated cargoes including proteins and nucleic acids have been regarded as biomarker for a variety of human diseases^17,18^. The lipid bilayer membrane of EVs provides exceptional stability to their molecular contents by safeguarding encapsulated nucleic acids from enzymatic degradation and temperature fluctuations that rapidly degrade free environmental DNA and RNA^19^. This protective capacity is particularly crucial in environmental applications where nucleic acids encounter harsh conditions such as temperature variations, enzymatic activity from nucleases and microbial metabolism, UV radiation, and pH fluctuations^20,21^.

EVs are actively released by living cells via regulated biogenesis processes that require ATP-dependent machinery, including VPS4 ATPase and ESCRT complexes, specific protein sorting mechanisms, and membrane rearrangement^15,22,23^. This active release process is fundamentally distinct from passive DNA release via cell lysis, apoptosis, or necrosis. This feature addresses a major limitation of conventional eDNA techniques: the inability to distinguish free DNA from living organisms versus dead cells, transported material, or past biological activity, leading to false-positive detections when aged or allochthonous DNA is present^24–26^.

Beyond nucleic acid preservation, EVs contain proteins that reflect organismal physiological state and health. Similar to environmental RNA’s capacity to detect stress-response genes and assess population health ^10^, EV-associated proteins provide functional information complementary to taxonomic identification. Protein biomarkers offer distinct advantages: greater stability than RNA under environmental stress^7^, post-translational modification information, and integration of cellular responses across multiple pathways^27^. While eRNA has demonstrated utility in detecting sublethal toxicity effects in fish through transcriptional biomarkers^6,10^,the multi-omic nature of EVs, containing both nucleic acids and proteins, enables simultaneous biodiversity assessment and functional health monitoring. This dual capacity represents a fundamental advantage over conventional eDNA/eRNA approaches that provide either taxonomic or functional information, but not both from a single sample.

Although EV-associated DNA has been used to characterize microbial communities in water^28,29^, its application to detecting macroorganisms has not been systematically evaluated. Here we develop and benchmark a workflow that recovers EV-encapsulated nucleic acids and proteins from water samples, and compare it directly against bulk eDNA and eRNA. All four methods (eDNA, eRNA, EV-DNA, EV-RNA) share an identical downstream workflow, extraction, amplification, 12S rRNA metabarcoding, and bioinformatic analysis, and differ only in the pre-analytical capture step: filtration for the bulk environmental fraction versus sequential 0.22 µm filtration, tangential flow filtration, and ultracentrifugation for the EV-encapsulated fraction. This design isolates the effect of the capture fraction itself rather than confounding it with differences in detection chemistry. We applied this comparison across two phases of increasing environmental complexity, a controlled aquarium of known fish species to assess detection accuracy and false-positive behaviour, and a field campaign at Xinglinwan Reservoir (XLW), Xiamen, China^30–32^, evaluating performance under real conditions, to test EV-associated nucleic acids and proteins as a complementary fraction for integrated biodiversity and physiological-signal monitoring.

## 2. Materials and Methods

### 2.1 Experimental Design

This study developed and benchmarked a workflow for recovering EV-encapsulated nucleic acids and proteins from water samples, comparing it directly against bulk eDNA and eRNA under standardized downstream conditions. A two-phase comparative design was used. Phase 1 was a controlled aquarium experiment with a known species composition to assess detection accuracy and false-positive behaviour. Phase 2 was field sampling at Xinglinwan Reservoir (XLW) to evaluate method applicability under real environmental conditions.

The core procedural sequence, nucleic-acid extraction, amplification, 12S rRNA metabarcoding, sequencing, and taxonomic assignment, is identical for all four methods. Only the pre-analytical capture step differs: 0.45 μm filter retention for bulk environmental nucleic acids (eDNA/eRNA), which captures the particle-bound and intra-membrane eDNA states^33^, versus sequential 0.22 μm filtration, tangential flow filtration, and ultracentrifugation of the downstream filtrate for the EV-encapsulated fraction (EV-DNA/EV-RNA)^28,29^. The recovery of intact, nucleic-acid-bearing EVs from natural waters by comparable workflows has been demonstrated in both aquatic microbial communities^34^ and macro-organism-derived aquatic eRNA^35^. EV-associated nucleic acids therefore represent a defined, membrane-bound subfraction of the total environmental nucleic-acid pool, one of several operationally separable eDNA states^36,37^, rather than a directly parallel target to bulk eNA, and the design isolates the effect of capture fraction from confounding differences in detection chemistry.

### 2.2 Sample Collection

Tank samples were collected from a 500 L recirculating commercial holding system in Xiamen, Jimei District, in which the seven study species shared a single connected water volume maintained at the supplier’s standard operating temperature. The tank had been stocked and operating in steady state for at least one week prior to sampling, providing an acclimation period over which EV release and free nucleic-acid shedding would equilibrate with the recirculating volume^38,39^. Five litres of water were collected as a single, well-mixed sample for downstream processing through the entire pipeline.

The seven species; *Oreochromis niloticus, O. mossambicus, Cyprinus carpio, Carassius carassius, Tachysurus fulvidraco, Anguilla japonica,* and *Channa argus,* were chosen for two reasons: they were the species commercially available in the local supplier’s system at the time of sampling, and all seven also occur in XLW based on prior regional surveys^32^. Validating the workflow first against a commercially available subset of XLW’s native ichthyofauna therefore provided a tractable known-composition mock community before deployment in the open field system.

All three field sites were sampled on the same day to minimise temporal variability. At each site, a single composite 10 L surface-water collection was processed through the workflow described in Section 2.3, with the resulting extract serving as one biological replicate per site. The controlled tank experiment was conducted as a single integrated sample per method, serving as a known-composition mock community for performance benchmarking rather than for population-level inference. Sampling volumes were chosen to ensure sufficient material for both bulk filter retention and downstream EV isolation, with the 5 L tank volume reflecting the closed, recirculating geometry of the holding system and the 10 L field volume reflecting the lower expected EV concentration and higher background turbidity of natural reservoir water.

Field sampling is performed in Xinglinwan Reservoir is located downstream of the Houxi River watershed, with an average depth of 2.5 m. Intensive urban growth, changes in agricultural land use, and sewage inputs have led to severe organic contamination, nutrient overloading, and diminished natural cleansing capacity; the reservoir continues to suffer from low dissolved oxygen levels and episodic fish-kill events^31^. Previous eDNA research at this site^32^ provided reference data for database construction. Sampling: Surface water samples (10 L each) were collected from three sites. Site X1 (coordinates: 24.601853°N, 118.064557°E); Site X2 (24.596585°N, 118.080861°E); Site X3 (24.581217°N, 118.082693°E). Weather conditions were sunny with no precipitation in the preceding 48 hours. All samples were stored at 4°C in sterile containers and processed within 12 hours of collection.

### 2.3 Fraction Isolation

Filtration methods and filter types show considerable diversity in eDNA studies. The most commonly used glass fiber filter membranes, such as GF/F and GF/A, vary based on size specifications. Other filter types include polycarbonate filters, cellulose nitrate, polyethylene sulfone, and polyvinylidene fluoride ^40,41^. Xiaoyu Chen et al. demonstrated that GF filters offer the highest detectability during eDNA analysis; accordingly, we used GF filters in this study^42^.

All four fractions (eDNA, eRNA, EV-DNA, EV-RNA) were derived from the same water sample at each site. First, 0.45 μm GF filters were used to capture large nucleic acid molecules, and these filters were designated for eDNA and eRNA isolation. The filtered water (filtrate) was retained for subsequent EV isolation steps. Each filter was divided in half: one half was used for DNA extraction (eDNA) and the other half for RNA extraction (eRNA), allowing paired nucleic acid analysis from the same sample. Each filter half was folded and placed into a spin-basket tube to maximize contact with lysis buffer during the incubation step.

To prepare the samples for EV separation, the remaining water was filtered through a 0.22 μm Nylon filter. This step removes large molecules and retains only EVs in the filtrate. Tangential flow filtration (TFF); TFF was performed to increase the EV concentration of the remaining water. This step continued until the water reached a suitable volume for ultracentrifugation (120-150 mL). Samples were processed through a Pellicon® 2 Mini Cassette Holder (MilliporeSigma) equipped with an Ultracel® 100 kDa membrane (0.1 m², Cat. No. P2C100C01). Ultracentrifugation; Samples from TFF were centrifuged using a Beckman L-80XP Ultracentrifuge at 100,000×g for 2 hours in No-Brake mode. The EVs that settled in the tubes were then resuspended with PBS (phosphate-buffered saline) for subsequent extraction. (Figure 1)

**Figure 1.**
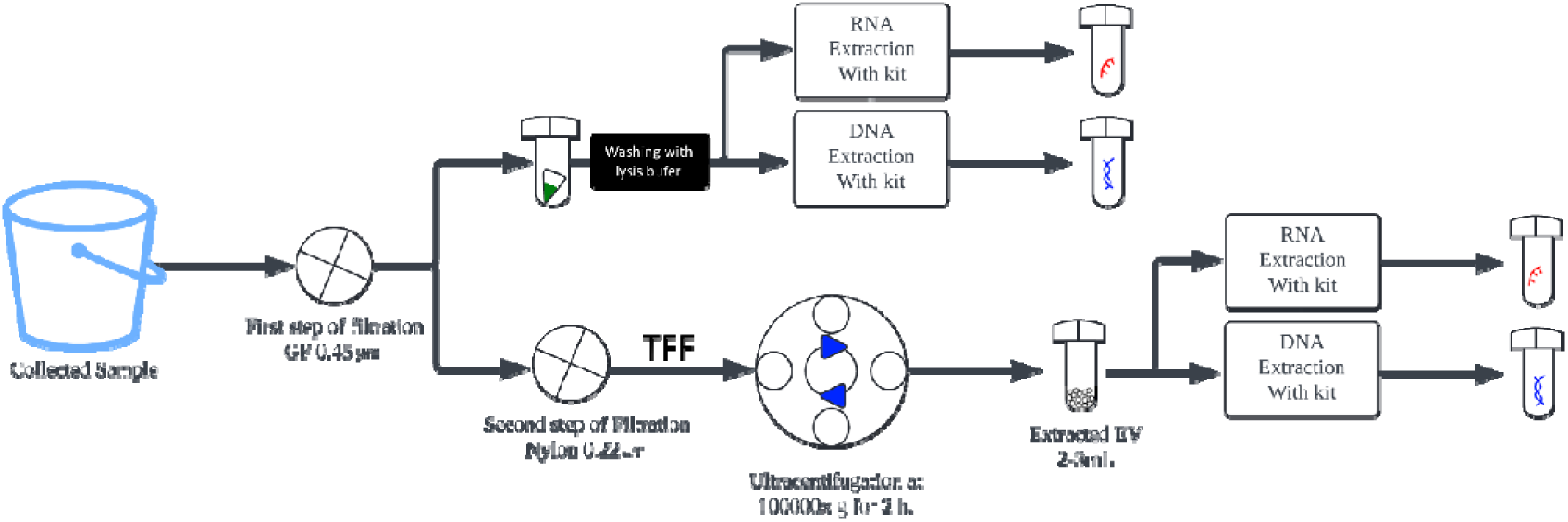
Main work scheme. First, water samples are collected and passed through a 0.45 µm GF filter. At this stage, a branching occurs: extraction is performed from the filter for eDNA/eRNA, while extraction is conducted directly from the water for EV-DNA/EV-RNA. For eNA, the filter is washed with lysis buffer, and extraction is carried out using a commercial kit. For EV-NA, after a subsequent 0.22 µm filtration, the sample undergoes 2 hours of ultracentrifugation at 100,000 x g, followed by the extraction from the liquid sample.

The four targets compared in this study are not strictly parallel by design, and this distinction is important for interpretation. Bulk eDNA and eRNA were recovered from material retained on the upstream 0.45 µm GF filter and therefore include intracellular DNA in trapped whole cells, debris-bound DNA, and any extracellular nucleic-acid aggregates above the filter cutoff. EV-DNA and EV-RNA were recovered from the downstream filtrate that passed through both the 0.45 µm and 0.22 µm filters and was then concentrated by tangential flow filtration and pelleted by ultracentrifugation. EV-NA therefore represents an operationally defined, membrane-bound subfraction of the total nucleic-acid pool, not a directly parallel measurement of the same material ^26^

Two consequences follow. First, the sub-0.22 µm filtrate retained for EV isolation is depleted of the larger particulate and cellular fractions that dominate bulk eDNA/eRNA, so the comparison is between distinct ecological capture fractions rather than between competing extraction chemistries applied to the same starting material. Second, residual free nucleic acid passing through 0.22 µm and surviving TFF/ultracentrifugation could in principle contribute to the EV-NA signal alongside genuinely vesicle-encapsulated material. We address this in two ways: (i) the 100 kDa TFF and 100,000×g ultracentrifugation steps enrich for membrane-bound vesicles at characteristic buoyant densities and sediment fractions^29,43^, conditions under which free, non-encapsulated nucleic acids and ribosomes are not efficiently retained; and (ii) the EV fraction was independently confirmed to contain intact, membrane-bound vesicles of the expected size range by Flow NanoAnalyzer particle counting and transmission electron microscopy (Figures S1, S2). Reported differences between EV-NA and bulk eNA therefore reflect differences between two operationally defined capture fractions, and any biological interpretation is framed accordingly throughout the manuscript.

### 2.4 The characterization of EVs

EV concentration and size distribution were measured for all isolated EV samples using a Flow NanoAnalyzer (NanoFCM Inc., Xiamen, China) calibrated against standard silica nanoparticles, with size determined from side-scatter intensity following the manufacturer’s protocol. EV morphology was assessed qualitatively by negative-stain transmission electron microscopy (TEM) using a Hitachi H-7650 microscope (Hitachi High-Technologies, Tokyo, Japan), which confirmed the typical cup-shaped vesicular morphology in the expected 30–200 nm size range (Figure S2).

### 2.5 Nucleic Acid Extraction

For DNA, we used the DNeasy Blood & Tissue Kit (QIAGEN, Germany). For both filter extraction and liquid EVs, samples were incubated with ATL buffer and Proteinase K for 1 hour at 56°C. For filters, samples were vortexed several times in a spin basket during the incubation period, then centrifuged at 12,000×g, and the protocol was continued with this residue, only on last step we used EB (10 mM Tris-Cl, pH 8.5) instead AE buffer which included in the kit. For EVs, the protocol was continued directly after incubation. DNA quality was assessed using a Qubit 4 fluorometer (Thermo Fisher Scientific, US). For RNA, the QIAGEN RNeasy Mini Kit was used. After addition of RLT buffer, samples were incubated at 37 °C for 15 min, and eRNA and EV-RNA were extracted according to the manufacturer’s protocol. Complementary DNA (cDNA) was generated using the HiScript II 1st Strand cDNA Synthesis Kit (Vazyme, Nanjing, China), primed with random hexamers to ensure unbiased capture of mRNA, rRNA, and small non-coding RNA species in the recovered transcript pool.

Additionally, it should be noted that during the extraction of eDNA and eRNA, high levels of PCR inhibitors were observed, which caused interference in the PCR in the initial stages, and to prevent this and obtain a sequenceable result, we used the OneStep PCR Inhibitor Removal Kit (Zymo Research, CA, USA).

### 2.6 Primer Selection and PCR Amplification

Candidate primers were screened in silico using AmplifX 2.1.1 (Nicolas Jullien; Aix-Marseille Université, CNRS, INP, Inst Neurophysiopathol, Marseille, France). Reference 12S sequences for fish species historically recorded in the study region were assembled, and candidate primers were evaluated for breadth of amplification across that list. Screening prioritised broad coverage of the regional ichthyofauna rather than fine-grained discriminatory power, because publicly available 12S reference sequences in this region do not reliably discriminate among some closely related cyprinid taxa (notably the *Hemiculter* / *Xenocypris* / *Pseudolaubuca* group); this taxonomic-resolution limit is acknowledged in the Discussion. Amplification targeted the hypervariable region of the mitochondrial 12S rRNA gene using MiFish primers^44^: forward 5′-GTYGGTAAAWCTCGTGCCAGC-3′and reverse 5′-CATAGTGGGGTATTAATCCYAGTTTG-3′. Degenerate positions (Y, W) were retained to broaden taxonomic coverage across divergent fish lineages. The MiFish primer set produced the strongest and most consistent in-silico amplification across the regional species list and was selected for all downstream PCR.

### 2.7 High-Throughput Sequencing

Library preparation and sequencing were performed on the Illumina NovaSeq 6000 platform using paired-end 250 bp (PE250) mode by MAGIGENE (Guangdong Magigene Technology Co., Ltd., Shenzhen, China). The MiFish 12S amplicon is approximately 170 bp, well within the read length of the PE250 platform, so paired-end reads fully overlap and merge to recover the complete amplicon. Library concentration was initially quantified using Qubit 4.0, and DNA library fragment integrity and insert size were verified using Qsep400. After quality control confirmation, libraries were pooled according to effective concentration and target data output requirements, loaded onto flowcells, and cluster generation was performed using cBOT. Samples were sequenced targeting the 12S rRNA region. Raw sequencing data were preprocessed to remove barcode sequences while retaining primer sequences, with forward primer reads assigned to R1 files and reverse primer reads to R2 files.

### 2.8 Bioinformatics Analysis

All metabarcoding analysis was performed using “eNA-Analyser” (https://github.com/jafarhayat/eNA-Analyser). Raw sequences were analyzed using the prepared reference database, and taxonomic results were obtained after metabarcoding.

A custom reference database was constructed by combining: (1) MiFish database sequences for regional fish species, (2) 12S rRNA sequences from NCBI GenBank. Detected species were validated against FishBase (www.fishbase.org) using a pre-specified regional-plausibility filter for subtropical Chinese freshwater systems applied uniformly to all four method outputs before downstream comparative analysis.

The pipeline was executed with default settings and with additional flags as “--min-identity 95” “--min-reads 5” for XLW and “--min-reads 10” for Tank sample, “--fish-split”, which were meant to; Quality Filtering Parameters: Paired-end reads underwent quality filtering via Trimmomatic (LEADING:3, TRAILING:3, SLIDINGWINDOW:4:20, MINLEN:50), followed by merging with FLASH (min overlap 10 bp, max 300 bp) and primer removal using Cutadapt (linked adapter syntax, error rate 0.15, min length 90 bp, max 250 bp). Different minimum read thresholds were applied based on sample type: 10 reads for tank samples (where known species composition allowed stricter filtering) and 5 reads for field samples (to maximize detection of rare species in the more complex natural community). Sensitivity analyses confirmed that results were robust to threshold selection within the 5-15 read range. BLAST Parameters: Taxonomic annotation used BLASTn against database with E-value ≤ 0.001, min percent identity = 95% (--min-identity 95), and min alignment length = 90 bp for confident species assignments.

### 2.9 Detection Classification and Performance Metrics

We compared eDNA, eRNA, EV-DNA, and EV-RNA detection using a pipeline that matches observed taxa to the local database by both exact name and BLAST similarity. Detections were filtered by abundance and confidence. Performance metrics, sensitivity, specificity, positive predictive value (PPV), and F1 score, were calculated from these classifications for each method and sampling site. This approach provides a robust, reproducible comparison of eNA and EV-NA detection accuracy and error rates. Alpha diversity metrics (species richness, Shannon index, Simpson index) were calculated using the vegan package v2.6-4. Reference species detection was assessed using unfiltered sequence data to maximize sensitivity for expected species.

### 2.10 Reference Species Detection

Reference species were defined differently by system. For the tank experiment, the reference set comprised the seven species physically stocked in the aquarium, constituting known ground truth. For the field site, the reference set comprised twelve species expected from prior biodiversity records; this expected list was treated as a benchmark for sensitivity rather than as exhaustive ground truth, acknowledging that the true assemblage may include additional taxa.

### 2.11 EV Protein Extraction and Metaproteomic Analysis

A subset of XLW EV samples collected across different sampling dates were analyzed by metaproteomics to validate biological origin and assess functional protein content. Sample preparation and mass spectrometry acquisition were performed as described in Xu et al.^34^. Briefly, EVs were collected at a single reservoir site across three sampling periods: spring (April), summer (August), and winter (December), yielding five, six, and seven samples respectively. The mass spectrometry metaproteomics data have been deposited to the ProteomeXchange Consortium (https://proteomecentral.proteomexchange.org) via the iProX partner repository^42,45^ with the dataset identifier PXD065377. Briefly, 100 μg of EV protein per sample was assessed by SDS-PAGE for quality verification, enzymatically digested with trypsin, and the resulting peptides were fractionated by high-pH reversed-phase UHPLC (Vanquish Flx, Thermo Scientific) using a gradient of 0–43% buffer B (80% acetonitrile) over 37 min. Equal quantities of fractionated peptides from each sample were combined and analyzed by LC-MS/MS on a Vanquish Neo system (Thermo Scientific) operating in data-dependent acquisition (DDA) mode, followed by data-independent acquisition (DIA) quantification.

For fish-targeted metaproteomic analysis in the present study, the DIA raw data were independently re-analyzed using DIA-NN version 2.3.2^46^. DIA-NN was operated in library-free mode, performing in silico digestion of the UniProt Swiss-Prot database^47^. Search parameters included: trypsin enzyme specificity with maximum 1 missed cleavage, peptide length 7–52 amino acids, precursor charge states 2–4, precursor m/z range 300–1800, mass accuracy 20 ppm (MS1 and MS2), oxidation of methionine as variable modification (maximum 2 per peptide), and 1% false discovery rate (FDR) at both precursor and protein group levels. Neural network classifiers and match-between-runs were enabled for optimal identification and quantification. Unique peptide sequences (≥7 amino acids) were extracted from DIA-NN output files (report.parquet) using custom Python scripts (Python 3.12, PyArrow 14.0, pandas 2.1). Taxonomic classification was performed using the Unipept Metaproteomics Analysis platform version 5.3^48,49^ with UniProt reference proteomes (2025.04 release), isoleucine/leucine equivalence enabled, and lowest common ancestor (LCA) algorithm for taxonomic assignment. Results were filtered to retain only fish taxa (class Actinopterygii). Taxonomic confidence scores ≥0.5 indicated reliable species-level assignments. Gene Ontology (GO) enrichment analysis was performed within Unipept to characterize molecular functions of fish-matched peptides.

For seasonal comparison; all samples were processed with the identical DIA-NN library-free workflow described in Section 2.11. To assign taxonomy and function at the peptide level, identified peptides were submitted to Unipept, and peptides confidently assigned to ray-finned fishes (Actinopterygii) were retained for the seasonal comparison. For each fish peptide, per-sample presence was determined from the DIA-NN precursor matrix and mapped to its sampling season. Because sequencing depth differed among seasons, functional representation was depth-normalized and expressed as peptides per 100 fish peptides detected within each season rather than as raw counts. Gene Ontology molecular-function terms were then tabulated per season to describe compositional differences in the fish EV proteome. Given the modest number of samples per season and the exploratory nature of this comparison, seasonal differences were treated as descriptive rather than statistically inferential, and functional categories represented by only a few peptides were flagged as insufficient for seasonal interpretation.

## 3. RESULTS

### 3.1 The performance metrics of EV-associated nucleic acids for biodiversity assessment in controlled Tank Experiment

EVs were successfully recovered from both the controlled tank and the XLW field samples. Mean particle concentrations measured by Flow NanoAnalyzer were 7.47 × 10⁸ particles/mL for tank samples and 2.37 × 10⁸ particles/mL for field samples (Figure S1), consistent with reported aquatic EV abundances. Transmission electron microscopy confirmed intact, cup-shaped membrane-bound vesicles of 30–200 nm diameter in all preparations (Figure S2), establishing that the downstream EV-DNA and EV-RNA analyses were performed on a morphologically and quantitatively validated EV fraction.

EV-DNA recovered all seven stocked reference species, the only target to do so. *Anguilla japonica* was detected by EV-DNA at low read depth (10 reads), placing this specific detection in a sensitivity-limited regime. In contrast, conventional eDNA, eRNA, and EV-RNA methods each detected six of seven reference species (85.7% sensitivity), failing to detect *Japanese eel* (Table 1, Figure 2A). Detection performance varied across methods when considering both sensitivity and precision (Figure 2B). While EV-DNA achieved perfect sensitivity (100%), it showed moderate precision (PPV = 53.8%) due to the detection of 6 additional non-reference fish species. Conventional eDNA showed the highest F1 score (0.75), balancing sensitivity (85.7%) with the highest precision (66.7%). EV-RNA demonstrated precision of 60.0% while detecting 10 total fish species (Figure 2C). The species detection breakdown revealed that EV-DNA detected the most reference species (7 true positives) but also the highest number of non-reference species (6 false positives), while conventional eDNA achieved the most favorable balance with 6 reference and only 3 non-reference detections. Detection efficiency was calculated as the number of species detected per 1,000 sequencing reads, enabling fair comparison between methods with different sequencing depths. EV-DNA demonstrated the highest detection efficiency at 3.01 species per 1,000 reads, approximately 1.9× higher than conventional eDNA (1.55 species/1,000 reads) (Figure 2D). EV-RNA showed intermediate efficiency (1.31 species/1,000 reads), while eRNA exhibited the lowest efficiency (1.12 species/1,000 reads).

**Figure 2.**
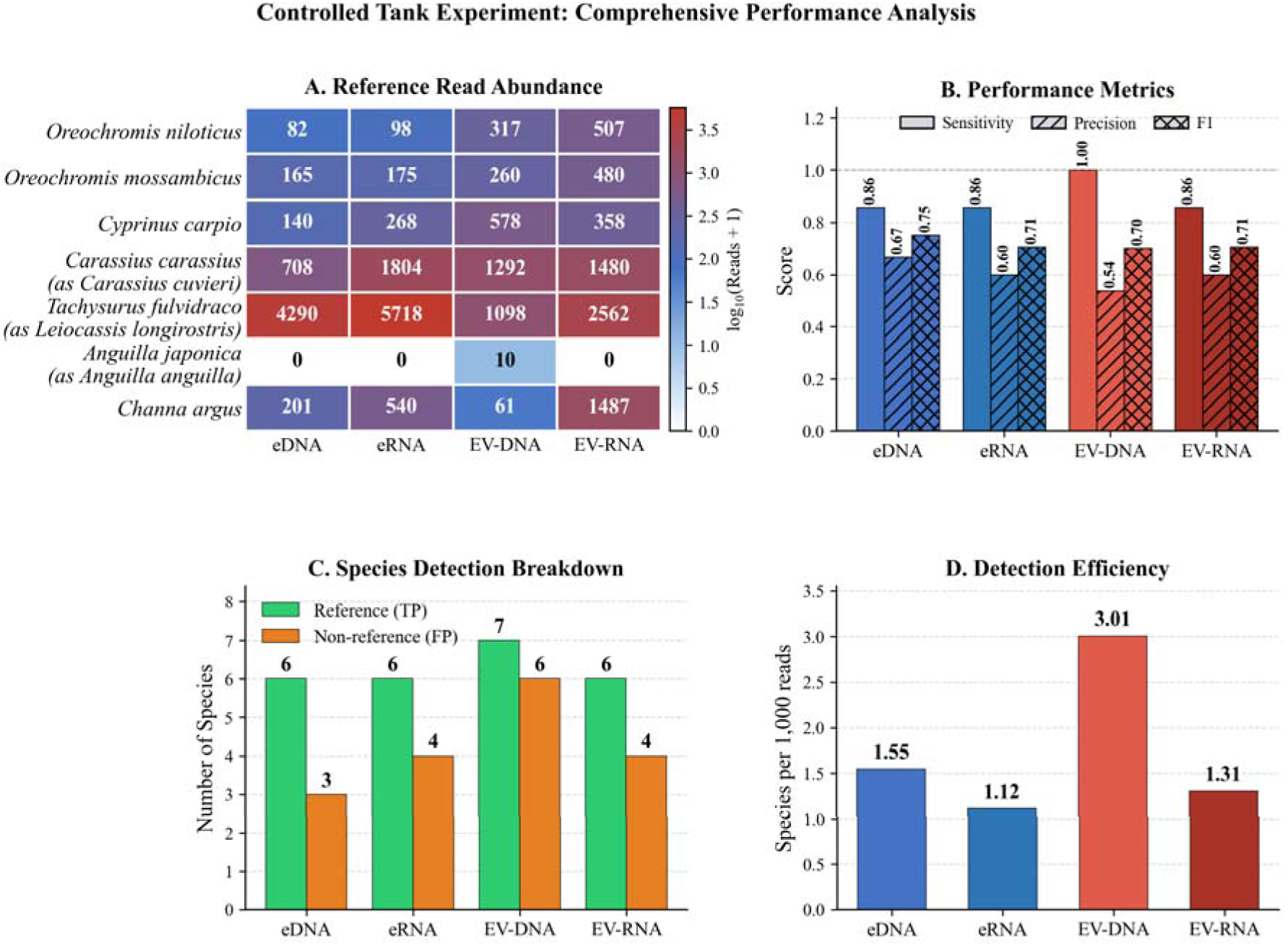
Comprehensive performance analysis of four molecular detection methods in the controlled tank experiment - (A) Read abundance heatmap showing sequencing read counts for seven reference fish species across all four methods. Cell values represent raw read counts; color intensity is scaled to log₁₀(Reads + 1). Anguilla japonica (Japanese eel) was detected exclusively by EV-DNA with 10 reads, while all other reference species were recovered by all methods. (B) Performance metrics including sensitivity (blue), precision/PPV (red), and F1 score (purple) for each method. EV-DNA achieved 100% sensitivity. (C) Species detection breakdown showing reference species (green, true positives) and non-reference species (orange, false positives) detected by each method. (D) Detection efficiency measured as species detected per 1,000 sequencing reads. EV-DNA showed the highest efficiency (3.01 species/1,000 reads), approximately 1.9× higher than conventional eDNA.

**Table 1.**
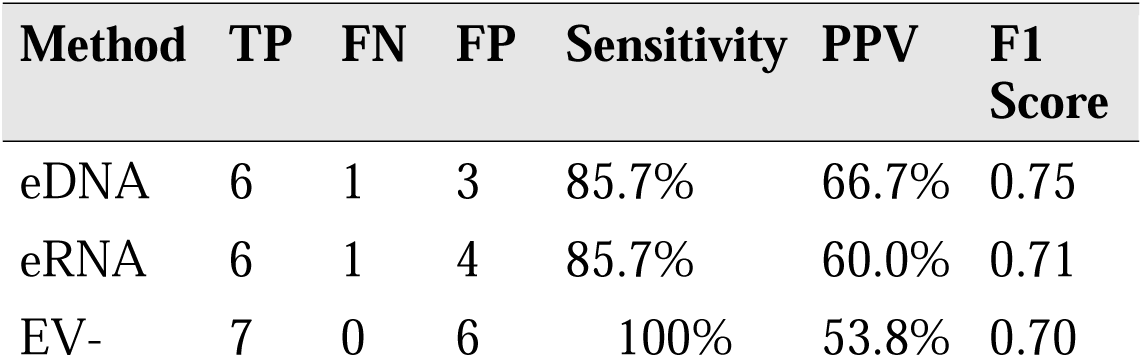
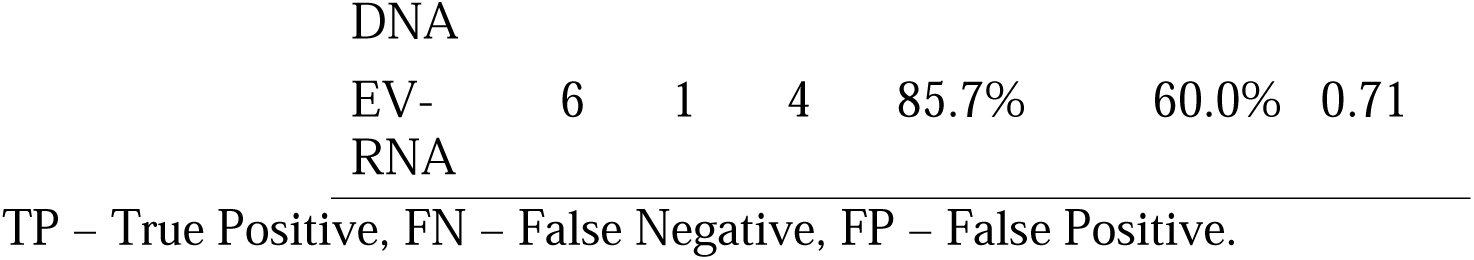
Detection performance metrics (eDNA, eRNA, EV-DNA, EV-RNA) in the controlled tank experiment (7 reference species)

| Method | TP | FN | FP | Sensitivity | PPV | F1 Score |
| --- | --- | --- | --- | --- | --- | --- |
| eDNA | 6 | 1 | 3 | 85.7% | 66.7% | 0.75 |
| eRNA | 6 | 1 | 4 | 85.7% | 60.0% | 0.71 |
| EV- | 7 | 0 | 6 | 100% | 53.8% | 0.70 |
| DNA |  |  |  |  |  |  |
| EV- | 6 | 1 | 4 | 85.7% | 60.0% | 0.71 |
| RNA |  |  |  |  |  |  |

TP – True Positive, FN – False Negative, FP – False Positive.

Read abundance for reference species varied considerably across methods (Figure 2A). *Tachysurus fulvidraco* (*Yellow catfish*) dominated read counts across all methods (1,098–5,718 reads), followed by *Carassius carassius* (708–1,804 reads). The EV-exclusive detection of *Anguilla japonica* by EV-DNA showed only 10 reads, demonstrating that EV methods can detect species at very low abundances that fall below the detection threshold of conventional approaches.

### 3.2 Field performance of EV-associated nucleic acids for biodiversity assessment at Xinglinwan Reservoir

Performance was evaluated against 12 species expected to occur at XLW based on the regional reference list. EV-RNA demonstrated substantially superior reference detection, successfully identifying 11 of 12 expected species (91.7% sensitivity). In contrast, eRNA detected 7 species (58.3% sensitivity), while both eDNA and EV-DNA each detected 6 expected species (50.0% sensitivity) (Table 2, Figure 3A–B). Notably, four expected species were detected exclusively by EV methods: *Cyprinus carpio* (*Common carp*; 94–136 reads), *Hemiculter bleekeri* (detected as *Xenocypris argentea*; 42–89 reads), *Hemiculter leucisculus* (detected as *Pseudolaubuca sinensis*; 15–43 reads), and *Monopterus albus* (*Asian swamp eel*; 15 reads, EV-RNA only). Only *Homatula laxiclathra* remained undetected by all methods.

**Figure 3.**
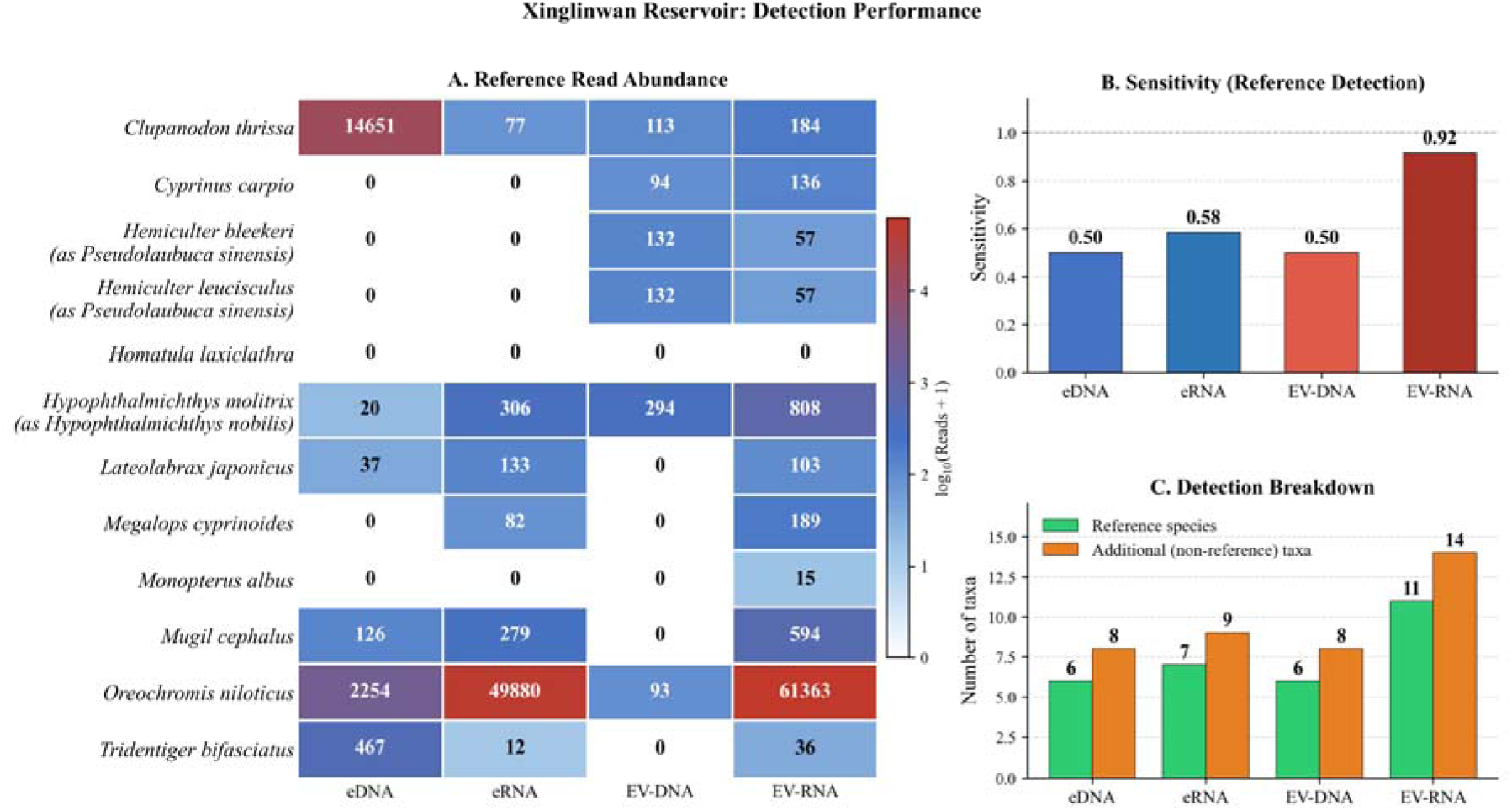
Comprehensive performance analysis of four molecular detection methods in XLW - A) Read abundance heatmap showing sequencing read counts for 12 expected species across all four methods. Cell values represent raw read counts; color intensity is scaled to log₁₀(Reads + 1). Four expected species were detected exclusively by EV-based methods: Cyprinus carpio, Hemiculter bleekeri (recovered as Xenocypris argentea), H. leucisculus (recovered as Pseudolaubuca sinensis), and Monopterus albus. Homatula laxiclathra was not detected by any method. (B) Performance metrics showing EV-RNA achieved highest sensitivity (0.92). (C) Species detection breakdown.

**Table 2.** Detection performance metrics (eDNA, eRNA, EV-DNA, EV-RNA) at Xinglinwan Reservoir (n = 12 expected species; n = 3 sites per method).

| Method | TP | FN | Sensitivity | PPV | F1 Score |
| --- | --- | --- | --- | --- | --- |
| eDNA | 6 | 6 | 50.0% | 42.9% | 0.46 |
| eRNA | 7 | 5 | 58.3% | 43.8% | 0.50 |
| EV-DNA | 6 | 6 | 50.0% | 42.9% | 0.46 |
| EV-RNA | 11 | 1 | 91.7% | 44.0% | 0.59 |

TP – reference species detected; FN – reference species not detected.

The species detection breakdown (Figure 3C) further illustrates the sensitivity–precision trade-off across methods. While EV-RNA detected the most expected species (11 of 12), it also recovered the largest number of additional, non-reference taxa (14). Conventional eDNA and EV-DNA each detected 6 expected species, and eRNA detected 7. Because the XLW expected-species list is a sensitivity benchmark rather than exhaustive ground truth, these additional taxa are treated as unverified detections rather than false positives, and precision-based metrics (PPV, F1) are reported only for the controlled tank experiment, where the species composition is known. The complete species detection matrix with binary detection status and raw read counts for all 25 taxa is provided in Table S4.

Read abundance for expected species revealed distinct patterns across methods (Figure 3A). *Oreochromis niloticus* dominated read counts, particularly in eRNA (49,880 reads) and EV-RNA (61,363 reads) samples. In contrast, the four EV-exclusive species showed moderate to low read counts (15–808 reads), indicating that EV methods successfully detected species present at lower abundances that were missed by conventional approaches. Also we need to note that, two expected species (*Hemiculter bleekeri* and *H. leucisculus*) were recovered under the names of closely related taxa (*Xenocypris argentea* and *Pseudolaubuca sinensis*).

Alpha diversity analysis revealed important differences in community representation across methods (Figure 4). While species richness showed high variability with no clear method-dependent pattern (Table S1), diversity indices differed substantially. EV-based methods showed markedly higher Shannon diversity (EV-DNA mean H’ = 1.24 ± 0.34; EV-RNA mean H’ = 0.95 ± 0.32) compared to conventional methods (eDNA mean H’ = 0.79 ± 0.59; eRNA mean H’ = 0.27 ± 0.38). Simpson diversity followed the same pattern, with EV-DNA showing the highest evenness (mean 1-D = 0.64 ± 0.07) compared to eRNA (mean 1-D = 0.11 ± 0.15). This pattern suggests that conventional methods captured communities dominated by a few highly abundant taxa (primarily Oreochromis spp.), while EV methods detected more evenly distributed communities with reduced dominance bias. Per-sample alpha diversity values for all indices are provided in Table S3.

**Figure 4.**
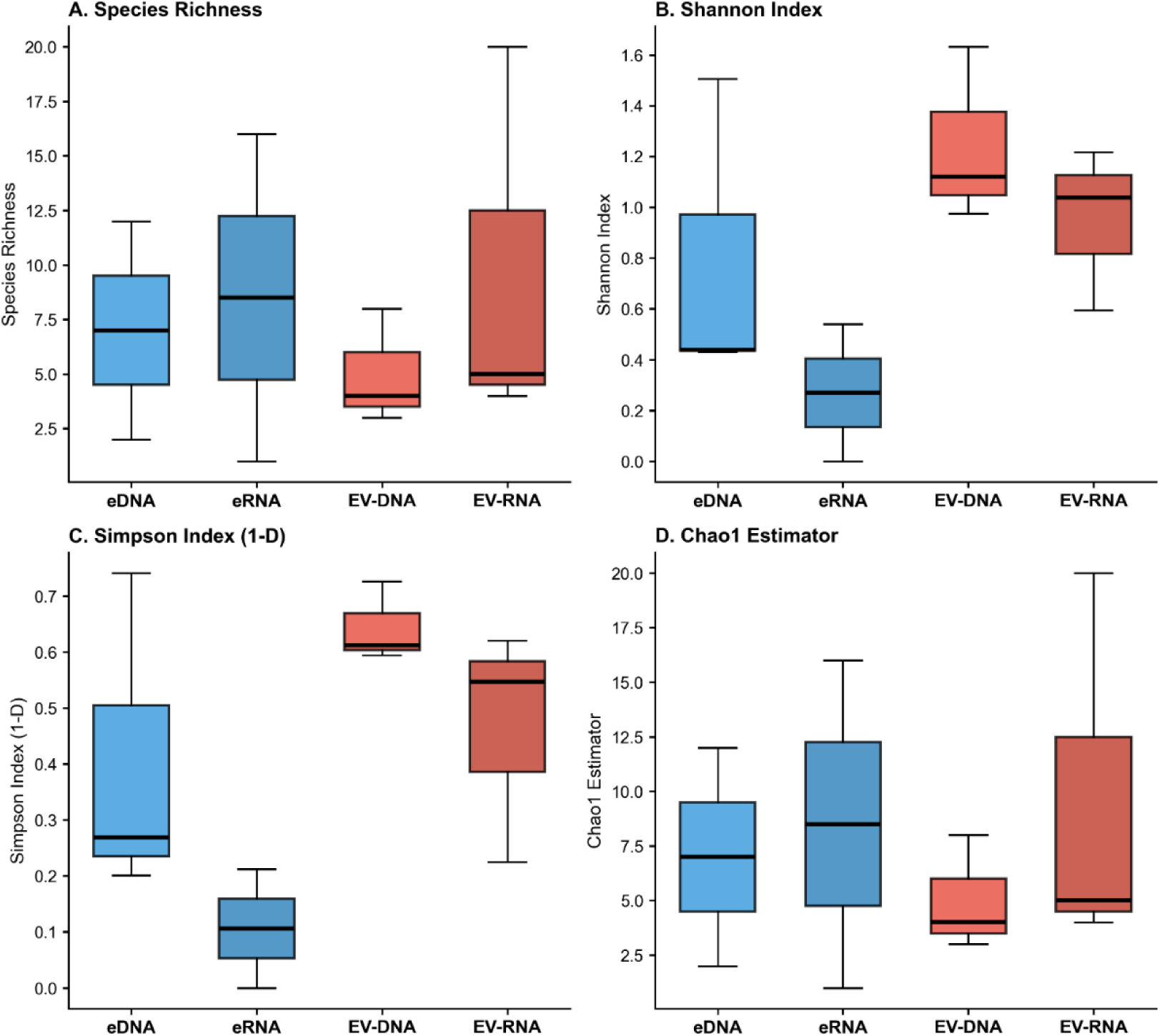
Alpha Diversity Comparison. Box plots comparing four alpha diversity metrics across detection methods: (A) Species richness, (B) Shannon diversity index, (C) Simpson diversity index (1-D), and (D) Chao1 richness estimator. Boxes show interquartile range with median line; whiskers extend to 1.5× IQR. Blue = conventional methods; red = EV-based methods.

Conventional eDNA showed the highest and most consistent human contamination (mean 86.8 ± 6.0%), followed by eRNA (mean 42.4 ± 36.3%) in field samples. EV-based methods showed lower mean contamination but higher variability: EV-DNA (mean 33.7 ± 56.6%) and EV-RNA (mean 25.6 ± 29.7%; individual sample values in Table S2). EV-RNA demonstrated 3.4-fold lower mean human contamination compared to eDNA, suggesting that EV isolation may partially exclude human-derived nucleic acids or that fish EVs were preferentially captured under the isolation conditions used.

### 3.3 Field EV metaproteomics confirms fish-derived biological origin

EVs from XLW was applied for metaproteomic analysis. Data-independent acquisition mass spectrometry generated 28,383–76,781 MS2 spectra per sample across EV samples analysed, indicating consistent analytical performance. DIA-NN analysis at 1% false discovery rate identified 163 fish protein groups across all samples, from which 1,453 unique peptide sequences (≥7 amino acids) were extracted for downstream taxonomic classification. Of these, 388 peptides (26.7%) were matched to proteins in the UniProt reference database, while 1,065 peptides (73.3%) remained unmatched (Figure 5A). Fish proteins from class Actinopterygii comprised the dominant matched taxonomic group, confirming that EVs isolated from lake water carry fish-derived biological material.

**Figure 5.**
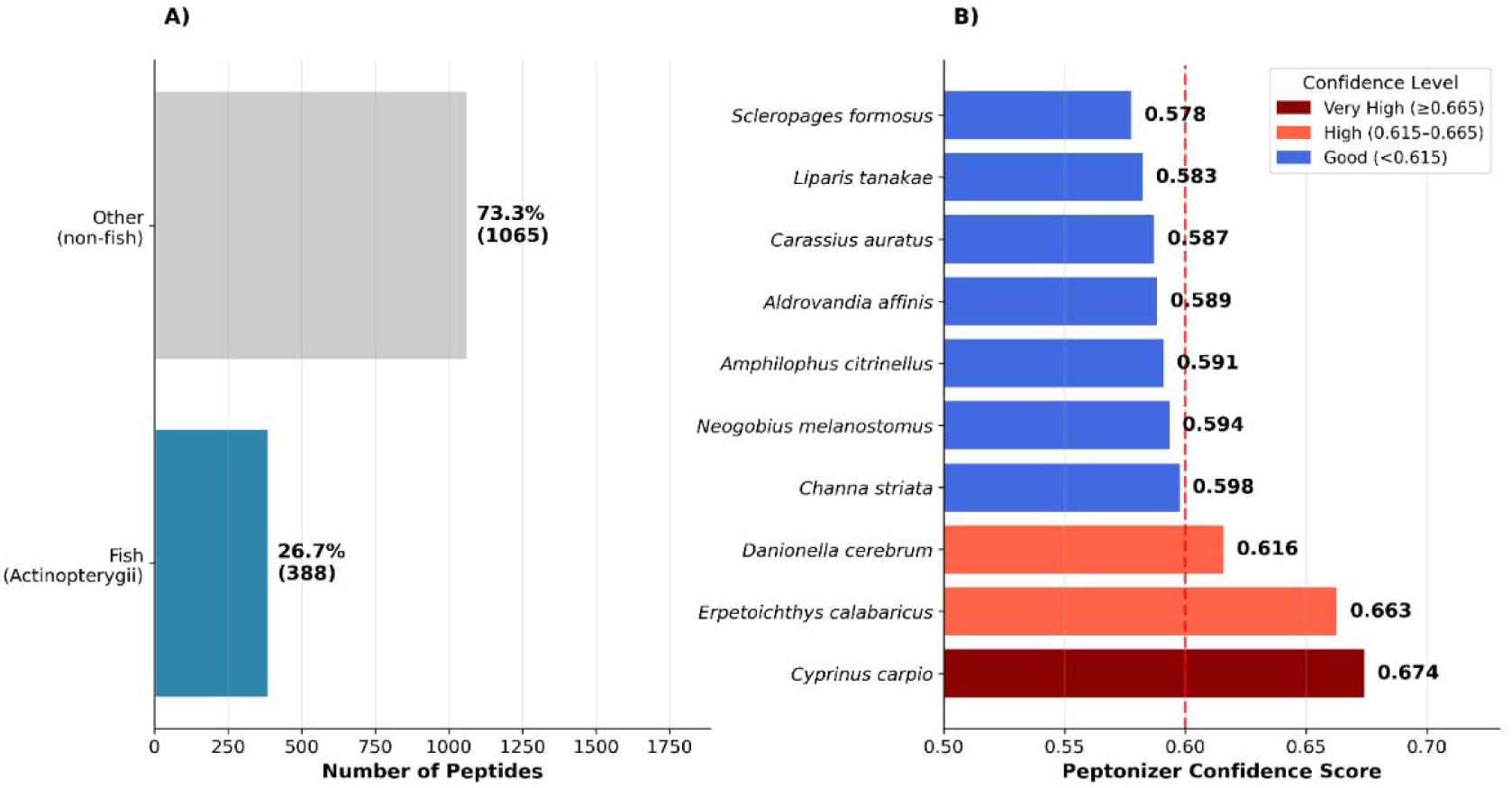
Metaproteomic detection of fish-derived proteins in lake water extracellular vesicles. (A) Taxonomic composition showing fish peptides (Actinopterygii, 26.7%) versus other taxa from 1,453 unique peptides analyzed. (B) Top 10 fish species detected with Unipept confidence scores.

Taxonomic classification using the lowest common ancestor algorithm identified proteins from 49 fish species spanning 28 orders and 44 families. As shown in Figure 5B, *Cyprinus carpio* exhibited the highest confidence score (0.674), indicating strong statistical support for the presence of this species’ proteins in XLW EVs. Notably, the proteomic detection of *Cyprinus carpio* independently corroborates its detection by EV-DNA and EV-RNA in the metabarcoding analysis, while this species was entirely absent from conventional eDNA and eRNA datasets.

### 3.4 Field EV-associated proteins show potential as fish physiological signals

Then we have predicted the function of EV-associated proteins. Gene ontology enrichment analysis of the 388 matched peptides identified 371 proteins (25.5%) with functional annotations across diverse molecular function categories. The most abundant categories were ATP binding (110 peptides, GO:0005524), GTP binding (62 peptides, GO:0005525), metal ion binding (56 peptides, GO:0046872), hydrolase activity (44 peptides, GO:0016787), and structural constituent of cytoskeleton (42 peptides, GO:0005200). Additional categories of direct relevance to fish physiological health assessment included oxidoreductase activity (14 peptides, GO:0016491), protein kinase activity (16 peptides, GO:0004672), zinc ion binding (22 peptides, GO:0008270), and translation elongation factor activity (18 peptides, GO:0003746). Full GO annotations with representative proteins and biological relevance are provided in Table 3.

**Table 3.**
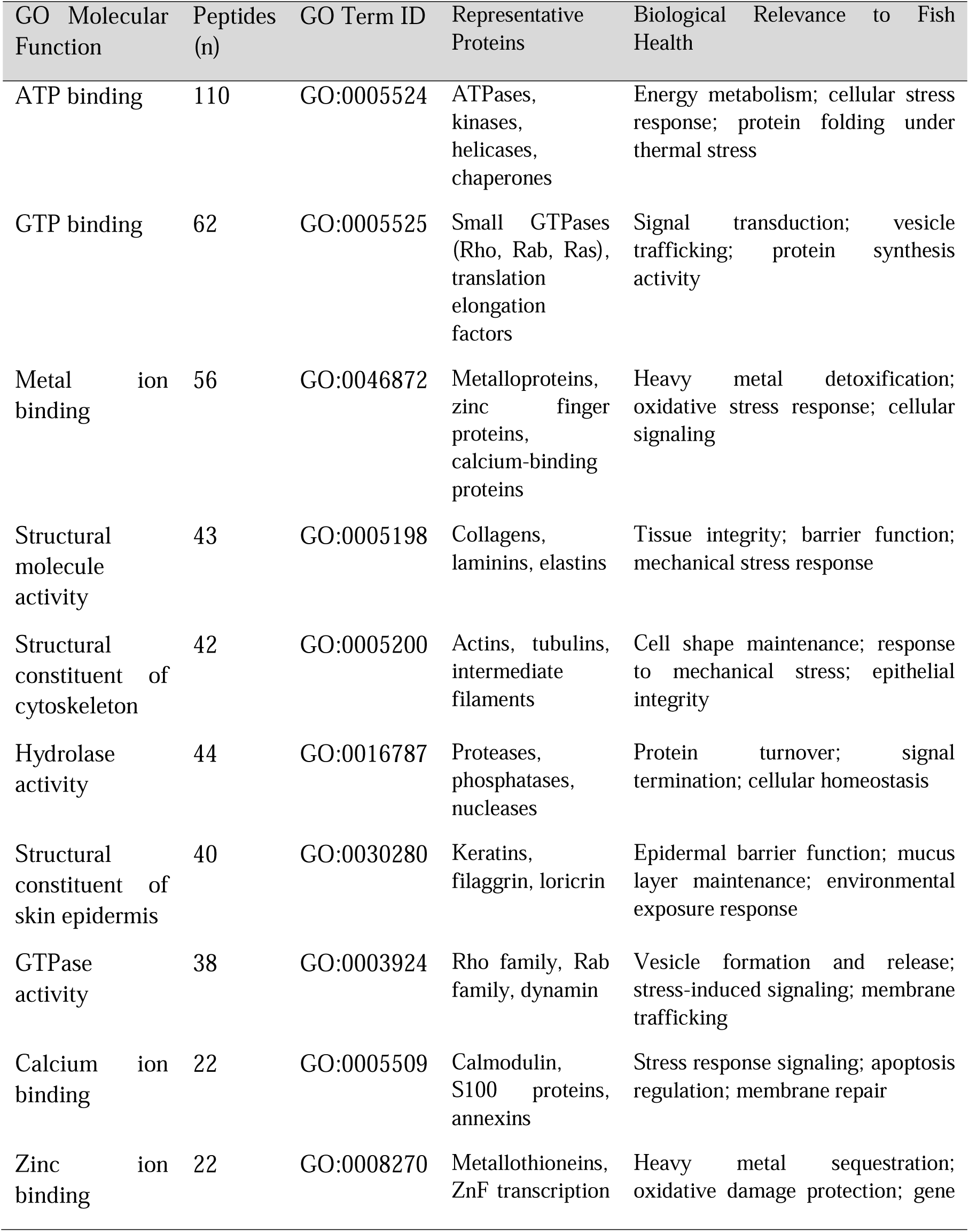
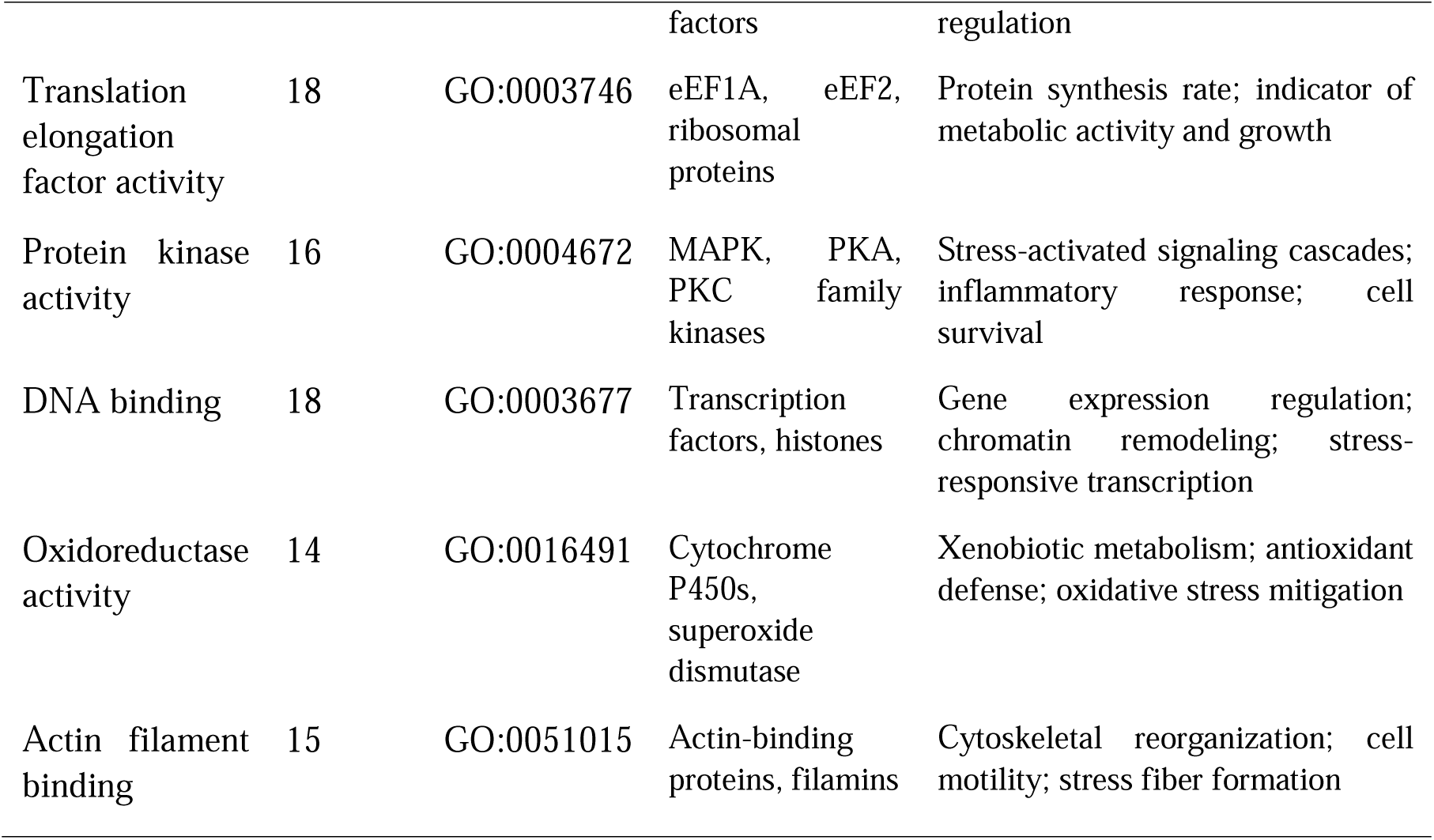
Predicted Functions of Fish Proteins Detected in XLW EVs.

Gene Ontology (GO) enrichment analysis identified 371 proteins with functional annotations from 388 matched peptides (26.7% of 1,453 unique peptides). Molecular functions shown represent top 15 categories by peptide abundance. Taxonomic confidence scores calculated within Unipept; functions with confidence ≥0.5 considered reliable. Representative proteins inferred from taxonomic assignments (primary match: *Cyprinus carpio*). Biological relevance based on established fish stress biomarker literature^27,50^

We then check whether these proteins change with environmental stresses. EVs from XLW were obtained across three seasons of a year, and then the relative abundance of signal proteins was compared (Figure 6). The bulk of the proteome was compositionally stable across seasons: the dominant molecular-function categories, including ATP binding, structural constituents, and hydrolase activity, showed comparable depth-normalized representation in all three periods. Against this stable background, two categories varied. GTP-binding and GTPase activity were markedly lower in summer than in spring or winter, and translation-elongation-factor activity was higher in winter. Categories corresponding to classical stress-associated functions (metal-ion binding, oxidoreductase and antioxidant activity) were detected but at peptide counts too low to support seasonal comparison (Supplementary Figure S6), and were therefore not interpreted quantitatively.

**Figure 6.**
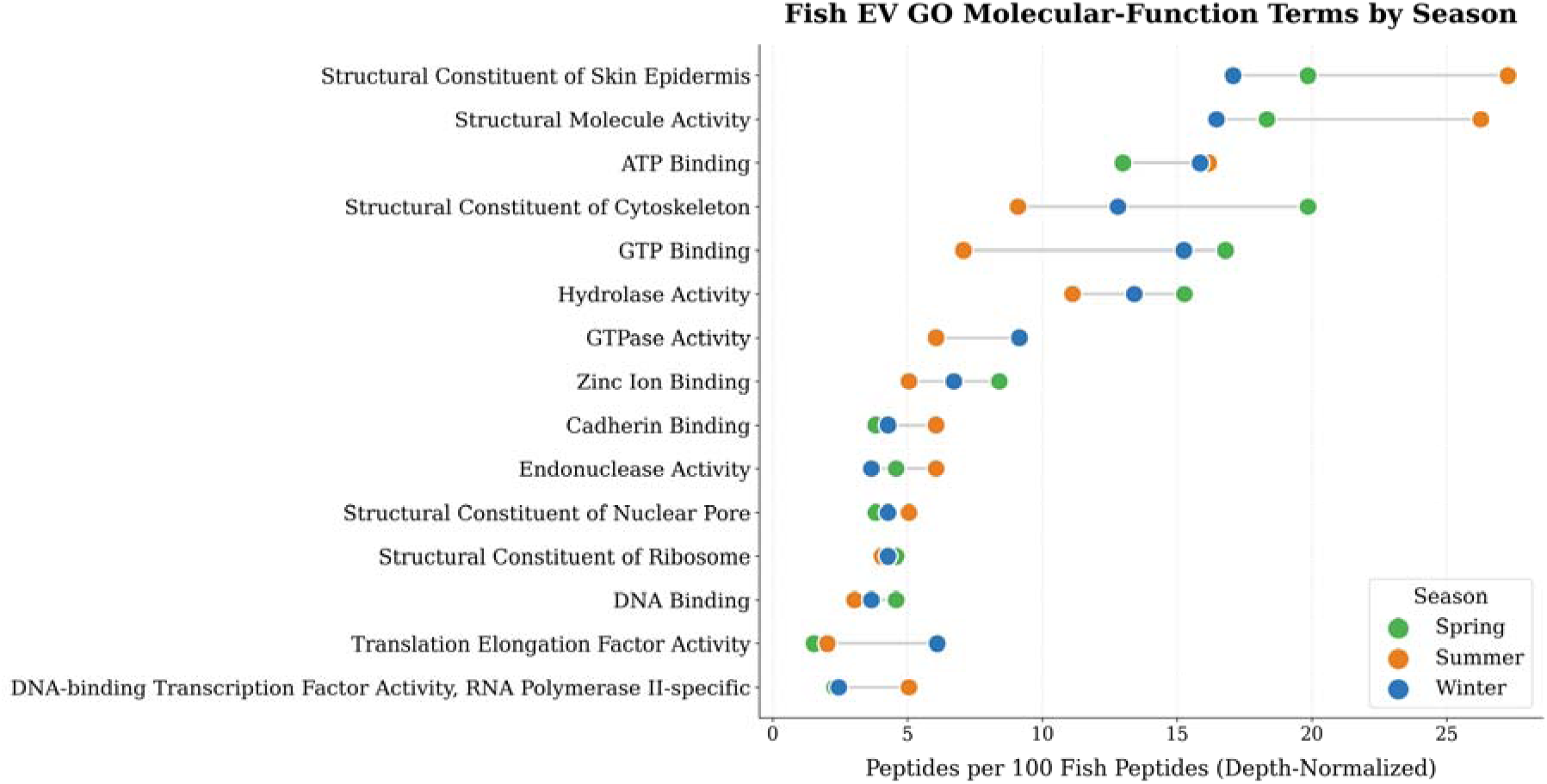
Seasonal representation of Gene Ontology (GO) molecular-function terms in the fish-derived extracellular-vesicle (EV) proteome at site He20, XLW. Each point is the mean depth-normalized abundance of peptides assigned to a given GO molecular-function category, expressed as peptides per 100 fish-derived peptides to correct for between-sample differences in identification depth, across three seasons: spring (April, n = 5; green), summer (August, n = 6; orange), and winter (December, n = 7; blue). The 15 categories (y-axis) are ordered by overall representation across seasons, from highest to lowest. GO category membership was assigned by keyword matching against protein descriptions and therefore represents a name-based proxy rather than a formal GO annotation; a single protein may contribute to more than one category.

## 4. DISCUSSION

This study provides a systematic evaluation of EV-associated nucleic acids as tools for fish biodiversity monitoring, comparing EV-DNA and EV-RNA directly against conventional eDNA and eRNA under both controlled and field conditions. The results demonstrate consistent advantages of EV-based approaches across control and field sampling contexts, supporting the hypothesis that membrane encapsulation of nucleic acids and proteins confers meaningful practical benefits for biomonitoring.

The superior sensitivity observed in EV-based methods, most notably EV-DNA achieved 100% sensitivity in the controlled aquarium experiment and EV-RNA achieved 91.7% sensitivity in the field. This could be attributed to the lipid bilayer membrane that physically shields encapsulated nucleic acids from environmental challenge. Free eDNA and eRNA in aquatic environments are subject to rapid breakdown by ubiquitous nucleases, with reported half-lives ranging from hours to days depending on temperature, UV exposure, pH, and microbial activity^51–53^. The EV membrane effectively extends the detection window for nucleic acids that would otherwise degrade below detection thresholds before sampling occurs. This mechanism is particularly relevant for species such as Anguilla japonica, Cyprinus carpio, and Monopterus albus, which were detected exclusively by EV methods in this study. These species may shed nucleic acids at lower rates, produce inherently less stable free nucleic acid signals, or occupy microhabitats that reduce their eDNA contribution to bulk water samples^38,39,54^. The broader community coverage of the EV-RNA fraction was reproduced at an independent urban water body (Yundang Lake), where EV-RNA again recovered more fish taxa than EV-DNA and re-detected several of the same species (Supplementary Table S6).

A further practical advantage of EV isolation emerged during sample processing. Substantial PCR inhibitor interference was encountered in eDNA and eRNA extracts collected from XLW, requiring use of a dedicated inhibitor removal kit before successful amplification could be achieved. No equivalent inhibition was observed in EV-based extracts. This pattern is consistent with the recognised challenge of PCR inhibitor co-extraction during filter-based eDNA collection from contaminated urban water bodies, where co-purified humic acids, heavy metals, and dissolved organic matter routinely suppress amplification efficiency^55^. The EV ultracentrifugation workflow involves pre-analytical separation from bulk water chemistry, which appears to substantially reduce inhibitor carryover. This represents an under-appreciated operational advantage of EV methods in polluted environments and warrants systematic evaluation in future protocol development studies.

An important finding is the divergence in performance between EV-DNA and EV-RNA across the two study systems. In the control aquarium experiment, EV-DNA outperformed all other methods in sensitivity, while in the field study, EV-RNA demonstrated markedly superior expected species recovery (91.7%) compared to EV-DNA (50.0%). This pattern may reflect differences in EV biogenesis under different physiological conditions. In a stable, controlled aquarium with well-fed fish at optimal temperature, cellular metabolism is regulated and DNA-containing EVs, likely derived from constitutive biogenesis pathways, may dominate. In contrast, fish in a stressed urban reservoir environment characterised by low dissolved oxygen, episodic fish kills, and organic contamination may preferentially release RNA-rich EVs as part of stress-response signaling.

This hypothesis is consistent with extensive biomedical evidence demonstrating that hypoxia markedly upregulates EV biogenesis through HIF-1α-dependent pathways and, in mammalian systems, enriches EV cargo with stress-response RNAs.^56,57^ Although this EV–hypoxia relationship has been characterised primarily in mammalian cell systems, the broad conservation of ESCRT machinery and HIF signaling pathways across vertebrates supports its plausibility in teleost fish under environmental stress^58^. If confirmed in aquatic organisms by future targeted studies, this would have important practical implications: EV-RNA may be the preferred method for field biomonitoring in contaminated environments, while EV-DNA may be more suitable for stable systems or controlled laboratory validation.

Beyond expected species detection, EV-based methods captured more evenly distributed communities as measured by Shannon and Simpson diversity indices. Conventional methods, particularly eRNA, showed strong dominance by *Oreochromis niloticus*, likely overwhelming the signal from less abundant co-occurring species. This pattern reflects a well-documented challenge in eDNA metabarcoding: A known limitation of eDNA metabarcoding is that dominant species can account for a disproportionate share of sequencing reads, reducing the likelihood of detecting less abundant taxa, a phenomenon termed ‘species masking’ ^59,60^. EV methods showed reduced dominance bias, with reads distributed more evenly across a broader taxonomic range. For biodiversity monitoring, this is a meaningful practical advantage, as it reduces the risk of common species masking the signals of rare or ecologically sensitive ones.

A recurring feature of the field detections was the recovery of two expected cyprinids, *Hemiculter bleekeri* and *H. leucisculus*, under the names of closely related taxa (*Xenocypris argentea* and *Pseudolaubuca sinensis*, respectively) and exclusively in the EV fractions. This mirrors the limited discriminatory power of publicly available 12S references within the *Hemiculter* / *Xenocypris* / *Pseudolaubuca* clade noted in the Methods. Both assigned species are nonetheless ecologically plausible for XLW: *X. argentea* is widely distributed across Chinese freshwater systems, and *P. sinensis* carries the historical synonym *Parapelecus fukiensis*, originally described from Fujian Province where the reservoir is located. We therefore propose two interpretations: either the detections represent cross-assignment of genuine *Hemiculter* signals due to reference-database limitations, or they reflect the actual presence of *Xenocypris* and *Pseudolaubuca* species that prior morphological surveys misidentified, a precedent explicitly documented within this clade^61^. In either case, the EV-exclusive nature of these detections supports our central conclusion regarding superior EV sensitivity for detecting low-abundance and closely related taxa.

Field EVs proteome provide a list of candidate protein markers that can indicate the The dominant functional category, ATP binding, encompasses the heat shock protein (HSP) family, whose upregulation is a well-established cellular response to thermal, osmotic, and chemical stressors in teleost fish ^62,63^. The co-occurrence of protein kinase activity is consistent with activation of stress-responsive signalling cascades, which are upstream regulators of HSP expression and anti-apoptotic pathways.

Particularly notable is the detection of metal ion-binding proteins and zinc ion-binding proteins, with metallothioneins explicitly identified among the zinc-binding category. Metallothioneins are tier-1 biomarkers for sublethal heavy metal exposure in fish, their expression is directly induced by cadmium, zinc, copper, and lead contamination, and their detection in environmental EVs provides a non-lethal, non-invasive signal of metal pollution that conventional eDNA methods cannot deliver^64^. Cytochrome P450 1A (CYP1A) is the primary enzymatic responder to polycyclic aromatic hydrocarbons and organochlorines in fish, while superoxide dismutase elevation indicates elevated reactive oxygen species consistent with xenobiotic exposure^65^. In addition, Rab GTPases regulate endosomal sorting and EV cargo loading, and their enrichment in the EV proteome indicates that the detected proteins were likely actively packaged into EVs through regulated biogenesis pathways rather than representing passive cell debris contamination^66^. The concurrent recovery of housekeeping and structural proteins (e.g., cytoskeletal constituents, actin-binding proteins) is itself expected for genuine EV cargo, which characteristically comprises both constitutive structural components and condition-responsive functional proteins; their presence therefore supports vesicular origin rather than indicating a specific physiological state.

The seasonal comparison of the fish EV proteome, though exploratory, revealed a pattern of broad compositional stability punctuated by a small number of functional shifts. The elevated representation of translation-elongation machinery in winter is consistent with the compensatory increase in protein-synthesis capacity that ectotherms are known to mount under cold acclimation, whereby low temperatures are offset by upregulation of translational components to maintain proteostasis.^62,67^ The lower representation of GTP-binding and GTPase functions in summer is more difficult to attribute definitively; because Rab and related small GTPases are central to extracellular-vesicle biogenesis and cargo loading^63,66^, seasonal variation in this category may reflect changes in vesicle production or packaging rather than a specific physiological state, and we interpret it cautiously. Importantly, the classical stress-associated proteins (metallothioneins, antioxidant enzymes, heat-shock proteins) were recovered at too few peptides in this dataset to support any seasonal claim; we therefore present them not as evidence of a seasonal stress response but as candidate targets that a more deeply sampled, temporally resolved study could pursue. Taken together, these observations indicate that environmental EV proteomes carry a detectable and partly time-varying physiological signal, and that, with denser sampling and orthogonal validation, EV-based metaproteomics has potential to track seasonal physiological change at the level of whole aquatic communities rather than individual organisms. Further study on controlled exposure conditions to build reference proteomes that link specific EV-protein signatures to defined physiological states would be required to provide basis for biomonitoring^68^. This dual capacity, simultaneous biodiversity assessment and recovery of candidate physiological-signal proteins from a single water sample, represents a meaningful expansion of what environmental nucleic acid methods can deliver^69,70^.

Several limitations of this study should be acknowledged. EV isolation by ultracentrifugation is more time-intensive and equipment-dependent than filtration-based eDNA collection, which may limit adoption in resource-constrained monitoring programmes. However, several alternative EV isolation approaches, including size-exclusion chromatography, polyethylene glycol precipitation, and membrane-affinity methods, have demonstrated comparable or superior yield and purity relative to ultracentrifugation while offering greater practical scalability.^43,71,72^ Commercial single-step EV isolation kits, now widely available, further lower the technical barrier: they require no ultracentrifuge and can be applied with standard benchtop equipment, making EV-based sampling tractable for smaller laboratories and routine monitoring programmes that lack specialised infrastructure^73–75^. Systematic optimisation of these protocols for environmental water matrices with high particulate loads, dissolved organic matter, and ionic complexity should be a priority for future work. The sample replication at XLW limits the statistical power of all comparative analyses. While the consistency of EV performance advantages across independent sites and the mechanistic interpretability of the findings provide confidence in the principal conclusions, all quantitative comparisons should be treated as effect estimates pending larger-scale validation in the future. Finally, the taxonomic discrepancies identified for two Cyprinidae species highlight the need for more comprehensive regional reference databases, particularly for species-rich families where 12S rRNA resolution is inherently limited.^76^

## 5. CONCLUSIONS

This study demonstrates that EV-associated nucleic acids recovered from environmental water samples can expand the detectable fraction of fish communities, including rare and low-abundance taxa potentially missed by conventional eDNA and eRNA approaches, while simultaneously providing access to candidate physiological-signal proteins and the signal change by seasons. Applied to impacted urban freshwater, where chemical contamination, eutrophication, and anthropogenic disturbance are ongoing, this multi-omic EV-based framework offers a non-invasive, single-sample strategy for integrated biomonitoring. The approach has potential relevance for regulatory ecological assessment programs seeking candidate physiological-signal indicators to complement species presence–absence data, though further optimisation of EV isolation protocols to reduce non-target nucleic acid contamination will be necessary for routine field deployment. Future work should prioritise method standardisation across water types, evaluating alternative EV extraction protocols, seasonal sampling to capture community dynamics, and validation against independent biodiversity surveys to establish detection thresholds suitable for policy-relevant monitoring frameworks.

## ASSOCIATED CONTENT

### Supporting Information

EV morphology and integrity assess ent, EV quantification by nano-flow cytometry and back-calculation of environmental concentrations, non-reference fish species detected by 4 methods at Xinglinwan Reservoir, Kruskal-Wallis test results for alpha diversity, human DNA contamination per sample, per-sample alpha diversity indices, complete species detection matrix with read counts.

### Data And Materials Availability

Raw sequencing data have been uploaded to Science Data Bank https://doi.org/10.57760/sciencedb.28036. All data needed to evaluate the conclusions in the paper are present in the paper and the Supporting Information.

## ACKNOWLEDGMENTS

This work was supported by Jing-Jin-Ji Regional Integrated Environmental Improvement-National Science and Technology Major Project (2026ZD1211600), Fujian Provincial Natural Science Foundation of China (2025J02030, 2025J01256), STS Project of Science and Technology Program of Fujian Province (2026T3006) and National Basic Science Data Center “Environment Health DataBase” (NO. NBSDC-DB-21).

## AUTHOR CONTRIBUTIONS

Q.H. and J.H. designed the study. J.H. and X.X. performed the experiments and collected field and tank samples. J.H. conducted the data analysis and prepared the manuscript. L.L. assisted in manuscript preparation, Y.W. and R.D.A.L. contributed to field sampling; A.O. and Z.P. assisted with EV isolation. All authors reviewed and approved the final version.

## COMPETING INTERESTS

The authors declare that they have no competing interests.

## Notes

### Competing Interest Statement

The authors have declared no competing interest.

