## Supplementary files for "Extracellular vesicle–associated nucleic acids and proteins as a fraction for fish biodiversity monitoring and physiological-signal recovery"

The authors declare no actual or potential competing financial interests.

**This file includes:**

3 figures and 6 tables

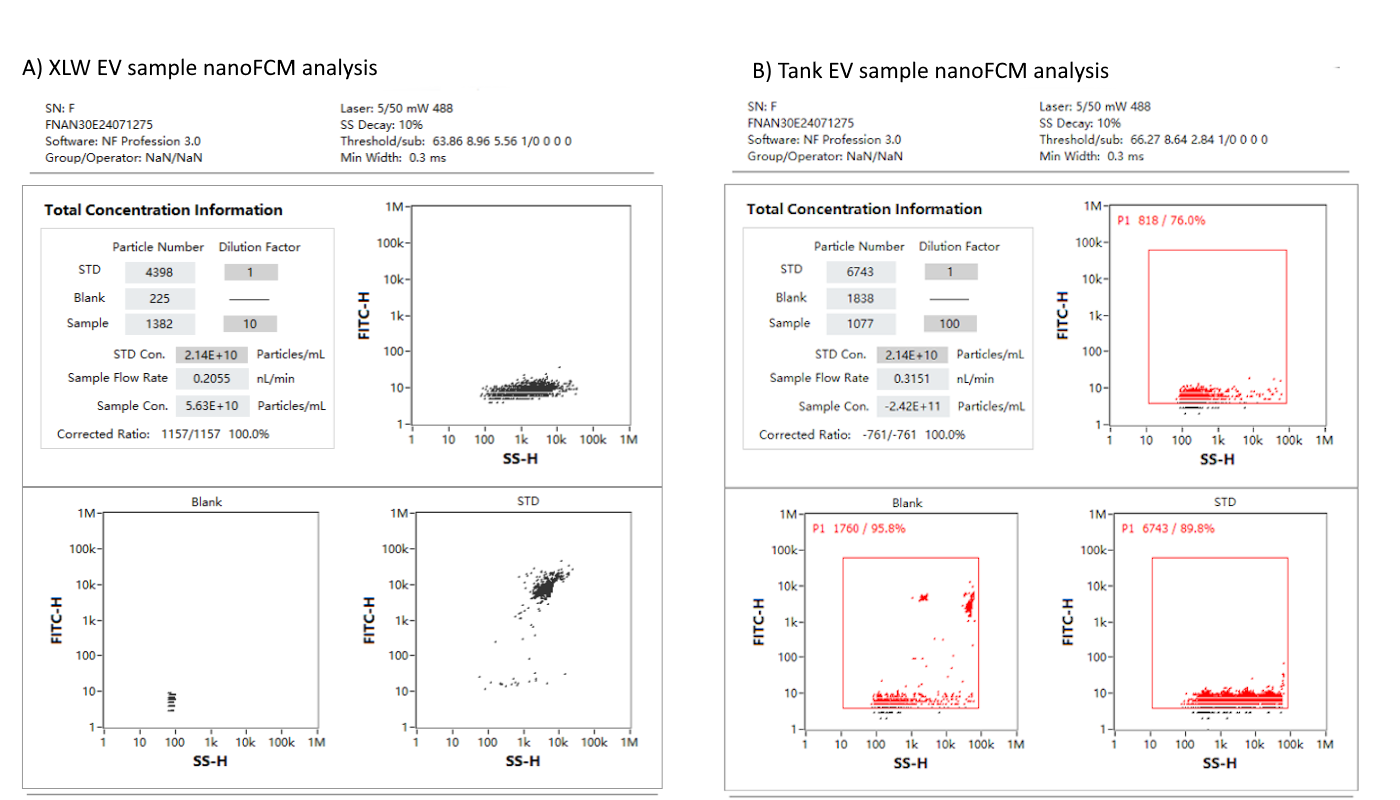

Figure S1. Representative nanoFCM output panels for EV samples from both study systems. (A) Xinglinwan Reservoir (XLW) EV sample (dilution factor 10): STD = 4,398 particles; blank = 225; sample = 1,382; instrument-reported concentration = 5.63×10^10^ particles/mL; flow rate = 0.2055 nL/min. (B) Controlled tank (T2) EV sample (dilution factor 100): STD = 6,743; blank = 1,838; sample = 1,077; flow rate = 0.3151 nL/min. Each panel shows the Total Concentration Information table (upper left) and scatter plots (FITC-H vs SS-H, logarithmic axes) for the sample, blank, and STD with the P1 nanoparticle detection gate (red box). EV concentration back-calculation: original environmental concentrations were derived as: (instrument reading × dilution factor) × eluate volume / source water volume. For XLW: 6.77×10^9^×100×3.5 mL/10,000 mL = 2.37×10^8^ particles/mL. For Tank: 3.36×10^12^ particles (total pelleted fraction)/4,500 mL = 7.47×10^8^ particles/mL.

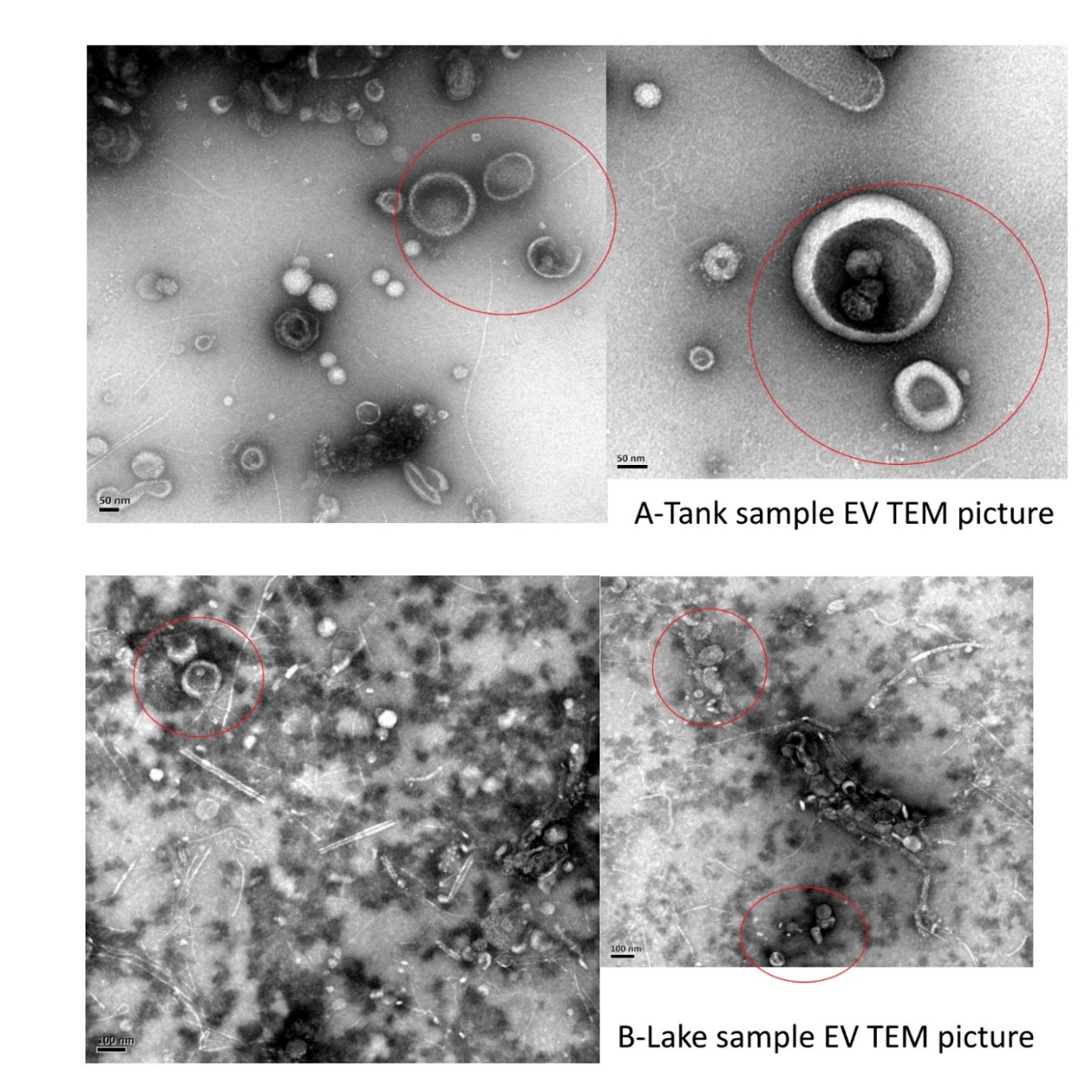

Figure S2. Transmission electron microscopy (TEM) images of isolated extracellular vesicles (EVs). EVs display the characteristic cup-shaped morphology and intact lipid bilayer membrane consistent with exosomes and microvesicles. Scale bars and imaging conditions as indicated. TEM analysis was performed on EV isolates prior to nucleic acid extraction to confirm vesicle integrity and absence of gross aggregation or membrane disruption.

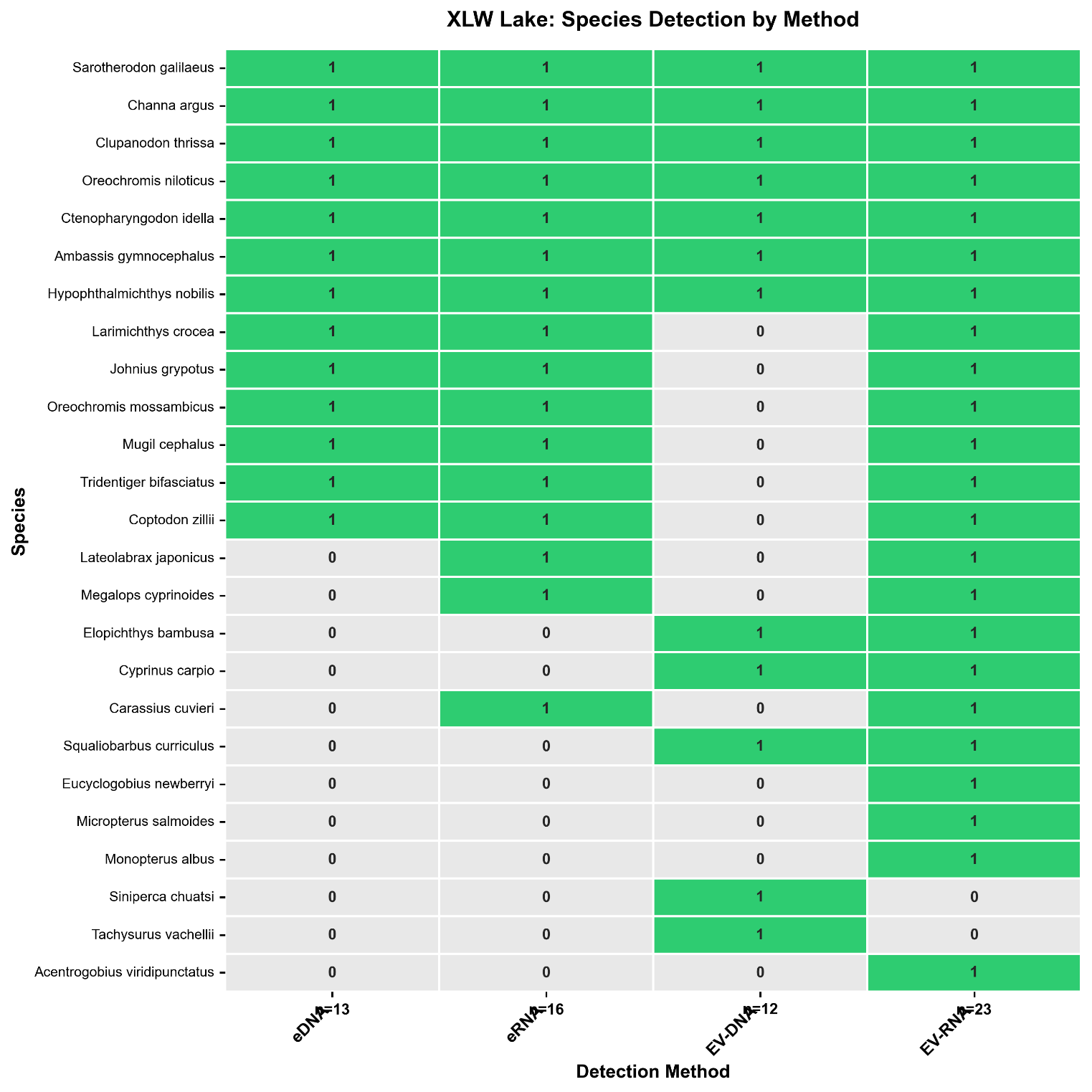

Figure S3. Non-reference fish species detected by ≥3 methods at Xinglinwan Reservoir (8 species). Green cells indicate detection; white cells indicate absence. Species detected by all four methods (Ambassis gymnocephalus, Channa argus, Ctenopharyngodon idella, Sarotherodon galilaeus) represent the highest-confidence ecological detections from non-reference taxa.

Table S1. Kruskal-Wallis test results for alpha diversity metrics across four detection methods at Xinglinwan Reservoir (XLW). n = 3 samples per method. No significant differences were detected for any metric (all p > 0.05).

| **Metric** | **H statistic** | **p-value** |
| --- | --- | --- |
| Richness | 0.609 | 0.894 |
| Shannon | 4.258 | 0.235 |
| Simpson | 4.424 | 0.219 |
| Chao1 | 0.609 | 0.894 |

Table S2. Human DNA contamination per sample at Xinglinwan Reservoir. Human reads were identified by alignment to the human reference genome (GRCh38) and excluded from downstream fish metabarcoding analysis.

| **Sample** | **Site** | **Method** | **Total reads** | **Human reads** | **Human (%)** |
| --- | --- | --- | --- | --- | --- |
| X1-eDNA | X1 | eDNA | 1,569 | 1,267 | 80.75 |
| X1-eRNA | X1 | eRNA | 476 | 324 | 68.07 |
| X1-EV-DNA | X1 | EV-DNA | 80,440 | 79,801 | 99.21 |
| X1-EV-RNA | X1 | EV-RNA | 637 | 373 | 58.56 |
| X2-eDNA | X2 | eDNA | 164,748 | 143,509 | 87.11 |
| X2-eRNA | X2 | eRNA | 92,006 | 15,398 | 16.74 |
| X2-EV-DNA | X2 | EV-DNA | 1,335 | 0 | 0.00 |
| X2-EV-RNA | X2 | EV-RNA | 130,928 | 24,001 | 18.33 |
| X3-eDNA | X3 | eDNA | 163,533 | 151,463 | 92.62 |
| X3-eRNA | X3 | eRNA | — | — | — |
| X3-EV-DNA | X3 | EV-DNA | 61,412 | 1,221 | 1.99 |
| X3-EV-RNA | X3 | EV-RNA | 1,503 | 0 | 0.00 |

Table S3. Alpha diversity indices per sample at Xinglinwan Reservoir. Richness = observed species; Shannon = H'; Simpson = 1-D; Chao1 = non-parametric richness estimator. Dash (—) indicates sample with no fish reads after quality filtering.

| **Sample** | **Site** | **Method** | **Richness** | **Total reads** | **Shannon** | **Simpson (1-D)** | **Chao1** |
| --- | --- | --- | --- | --- | --- | --- | --- |
| X1-eDNA | X1 | eDNA | 2 | 226 | 0.439 | 0.269 | 2.0 |
| X1-eRNA | X1 | eRNA | 1 | 36 | 0.000 | 0.000 | 1.0 |
| X1-EV-DNA | X1 | EV-DNA | 3 | 83 | 0.975 | 0.594 | 3.0 |
| X1-EV-RNA | X1 | EV-RNA | 4 | 93 | 1.037 | 0.547 | 4.0 |
| X2-eDNA | X2 | eDNA | 12 | 16,160 | 1.506 | 0.741 | 12.0 |
| X2-eRNA | X2 | eRNA | 16 | 56,313 | 0.540 | 0.213 | 16.0 |
| X2-EV-DNA | X2 | EV-DNA | 8 | 623 | 1.632 | 0.726 | 8.0 |
| X2-EV-RNA | X2 | EV-RNA | 20 | 69,779 | 0.594 | 0.225 | 20.0 |
| X3-eDNA | X3 | eDNA | 7 | 10,437 | 0.431 | 0.201 | 7.0 |
| X3-eRNA | X3 | eRNA | — | — | — | — | — |
| X3-EV-DNA | X3 | EV-DNA | 4 | 156 | 1.120 | 0.612 | 4.0 |
| X3-EV-RNA | X3 | EV-RNA | 5 | 872 | 1.216 | 0.621 | 5.0 |

Table S4. Complete species detection matrix with read counts for all 25 fish taxa detected at Xinglinwan Reservoir. Detection indicated as 1 (detected) or 0 (not detected). Reference species (those expected from prior surveys) are denoted in Table S4 of the species matrix file.

| **Species** | **eDNA** | **eDNA reads** | **eRNA** | **eRNA reads** | **EV-DNA** | **EV-DNA reads** | **EV-RNA** | **EV-RNA reads** | **# Methods** |
| --- | --- | --- | --- | --- | --- | --- | --- | --- | --- |
| Acentrogobius viridipunctatus | 0 | 0 | 0 | 0 | 0 | 0 | 1 | 10 | 1 |
| Ambassis gymnocephalus | 1 | 51 | 1 | 71 | 1 | 20 | 1 | 142 | 4 |
| Carassius cuvieri | 0 | 0 | 1 | 204 | 0 | 0 | 1 | 40 | 2 |
| Channa argus | 1 | 28 | 1 | 83 | 1 | 11 | 1 | 251 | 4 |
| Clupanodon thrissa | 1 | 14,651 | 1 | 77 | 1 | 113 | 1 | 184 | 4 |
| Coptodon zillii | 1 | 2,770 | 1 | 1,211 | 0 | 0 | 1 | 2,063 | 3 |
| Ctenopharyngodon idella | 1 | 15 | 1 | 48 | 1 | 55 | 1 | 292 | 4 |
| Cyprinus carpio | 0 | 0 | 0 | 0 | 1 | 94 | 1 | 136 | 2 |
| Elopichthys bambusa | 0 | 0 | 0 | 0 | 1 | 68 | 1 | 63 | 2 |
| Eucyclogobius newberryi | 0 | 0 | 0 | 0 | 0 | 0 | 1 | 49 | 1 |
| Hypophthalmichthys nobilis | 1 | 20 | 1 | 306 | 1 | 294 | 1 | 808 | 4 |
| Johnius grypotus | 1 | 10 | 1 | 14 | 0 | 0 | 1 | 43 | 3 |
| Larimichthys crocea | 1 | 9 | 1 | 14 | 0 | 0 | 1 | 108 | 3 |
| Lateolabrax japonicus | 0 | 0 | 1 | 76 | 0 | 0 | 1 | 28 | 2 |
| Megalops cyprinoides | 0 | 0 | 1 | 82 | 0 | 0 | 1 | 189 | 2 |
| Micropterus salmoides | 0 | 0 | 0 | 0 | 0 | 0 | 1 | 16 | 1 |
| Monopterus albus | 0 | 0 | 0 | 0 | 0 | 0 | 1 | 15 | 1 |
| Mugil cephalus | 1 | 15 | 1 | 29 | 0 | 0 | 1 | 97 | 3 |
| Oreochromis mossambicus | 1 | 275 | 1 | 3,070 | 0 | 0 | 1 | 2,390 | 3 |
| Oreochromis niloticus | 1 | 2,254 | 1 | 49,880 | 1 | 93 | 1 | 61,363 | 4 |
| Sarotherodon galilaeus | 1 | 6,258 | 1 | 1,172 | 1 | 12 | 1 | 2,394 | 4 |
| Siniperca chuatsi | 0 | 0 | 0 | 0 | 1 | 48 | 0 | 0 | 1 |
| Squaliobarbus curriculus | 0 | 0 | 0 | 0 | 1 | 26 | 1 | 27 | 2 |
| Tachysurus vachellii | 0 | 0 | 0 | 0 | 1 | 28 | 0 | 0 | 1 |
| Tridentiger bifasciatus | 1 | 467 | 1 | 12 | 0 | 0 | 1 | 36 | 3 |

Table S5 Proteins by season table:

| GO_MF | Spring | Summer | Winter | Spring_n100 | Summer_n100 | Winter_n100 | total |
| --- | --- | --- | --- | --- | --- | --- | --- |
| structural constituent of skin epidermis | 26 | 27 | 28 | 19.85 | 27.27 | 17.07 | 81 |
| structural molecule activity | 24 | 26 | 27 | 18.32 | 26.26 | 16.46 | 77 |
| ATP binding | 17 | 16 | 26 | 12.98 | 16.16 | 15.85 | 59 |
| structural constituent of cytoskeleton | 26 | 9 | 21 | 19.85 | 9.09 | 12.8 | 56 |
| GTP binding | 22 | 7 | 25 | 16.79 | 7.07 | 15.24 | 54 |
| hydrolase activity | 20 | 11 | 22 | 15.27 | 11.11 | 13.41 | 53 |
| GTPase activity | 12 | 6 | 15 | 9.16 | 6.06 | 9.15 | 33 |
| zinc ion binding | 11 | 5 | 11 | 8.4 | 5.05 | 6.71 | 27 |
| cadherin binding | 5 | 6 | 7 | 3.82 | 6.06 | 4.27 | 18 |
| endonuclease activity | 6 | 6 | 6 | 4.58 | 6.06 | 3.66 | 18 |
| structural constituent of nuclear pore | 5 | 5 | 7 | 3.82 | 5.05 | 4.27 | 17 |
| structural constituent of ribosome | 6 | 4 | 7 | 4.58 | 4.04 | 4.27 | 17 |
| DNA binding | 6 | 3 | 6 | 4.58 | 3.03 | 3.66 | 15 |
| translation elongation factor activity | 2 | 2 | 10 | 1.53 | 2.02 | 6.1 | 14 |
| DNA-binding transcription factor activity, RNA polymerase II-specific | 3 | 5 | 4 | 2.29 | 5.05 | 2.44 | 12 |
| DNA nuclease activity | 4 | 4 | 4 | 3.05 | 4.04 | 2.44 | 12 |
| nuclear receptor binding | 3 | 4 | 4 | 2.29 | 4.04 | 2.44 | 11 |
| transcription coactivator activity | 3 | 4 | 4 | 2.29 | 4.04 | 2.44 | 11 |
| ubiquitin protein ligase binding | 4 | 3 | 3 | 3.05 | 3.03 | 1.83 | 10 |
| catalytic activity | 3 | 3 | 4 | 2.29 | 3.03 | 2.44 | 10 |
| RNA polymerase II cis-regulatory region sequence-specific DNA binding | 3 | 4 | 3 | 2.29 | 4.04 | 1.83 | 10 |
| protein tag activity | 3 | 3 | 3 | 2.29 | 3.03 | 1.83 | 9 |
| metal ion binding | 2 | 2 | 4 | 1.53 | 2.02 | 2.44 | 8 |
| nucleic acid binding | 3 | 2 | 3 | 2.29 | 2.02 | 1.83 | 8 |
| methyltransferase activity | 2 | 2 | 3 | 1.53 | 2.02 | 1.83 | 7 |
| structural constituent of postsynaptic actin cytoskeleton | 2 | 2 | 3 | 1.53 | 2.02 | 1.83 | 7 |
| transmembrane signaling receptor activity | 2 | 2 | 2 | 1.53 | 2.02 | 1.22 | 6 |
| G protein-coupled receptor activity | 2 | 2 | 2 | 1.53 | 2.02 | 1.22 | 6 |
| actin binding | 3 | 1 | 2 | 2.29 | 1.01 | 1.22 | 6 |
| protein heterodimerization activity | 3 | 0 | 3 | 2.29 | 0 | 1.83 | 6 |
| structural constituent of chromatin | 3 | 0 | 3 | 2.29 | 0 | 1.83 | 6 |
| calcium ion binding | 1 | 1 | 4 | 0.76 | 1.01 | 2.44 | 6 |
| ATP-dependent protein folding chaperone | 1 | 1 | 3 | 0.76 | 1.01 | 1.83 | 5 |
| lipid binding | 2 | 1 | 2 | 1.53 | 1.01 | 1.22 | 5 |
| oxoglutarate-dependent dioxygenase activity | 1 | 1 | 2 | 0.76 | 1.01 | 1.22 | 4 |
| phosphatidylinositol-3-phosphate phosphatase activity | 2 | 0 | 2 | 1.53 | 0 | 1.22 | 4 |
| GTPase activator activity | 1 | 1 | 2 | 0.76 | 1.01 | 1.22 | 4 |
| cysteine-type deubiquitinase activity | 2 | 0 | 2 | 1.53 | 0 | 1.22 | 4 |
| ubiquitin binding | 2 | 0 | 2 | 1.53 | 0 | 1.22 | 4 |
| actin filament binding | 2 | 1 | 1 | 1.53 | 1.01 | 0.61 | 4 |
| galactosylceramide sulfotransferase activity | 1 | 1 | 1 | 0.76 | 1.01 | 0.61 | 3 |
| RNA binding | 1 | 0 | 2 | 0.76 | 0 | 1.22 | 3 |
| metallopeptidase activity | 1 | 1 | 1 | 0.76 | 1.01 | 0.61 | 3 |
| cytoskeletal motor activity | 1 | 1 | 1 | 0.76 | 1.01 | 0.61 | 3 |
| SNAP receptor activity | 1 | 1 | 1 | 0.76 | 1.01 | 0.61 | 3 |
| extracellularly ATP-gated monoatomic cation channel activity | 1 | 1 | 1 | 0.76 | 1.01 | 0.61 | 3 |
| oxidoreductase activity | 1 | 1 | 1 | 0.76 | 1.01 | 0.61 | 3 |
| protein serine/threonine kinase activity | 1 | 1 | 1 | 0.76 | 1.01 | 0.61 | 3 |
| purinergic nucleotide receptor activity | 1 | 1 | 1 | 0.76 | 1.01 | 0.61 | 3 |
| retinal binding | 1 | 0 | 2 | 0.76 | 0 | 1.22 | 3 |
| sphingosine N-acyltransferase activity | 1 | 1 | 1 | 0.76 | 1.01 | 0.61 | 3 |
| retinol binding | 1 | 0 | 2 | 0.76 | 0 | 1.22 | 3 |
| iron ion binding | 1 | 1 | 1 | 0.76 | 1.01 | 0.61 | 3 |
| steroid hydroxylase activity | 1 | 1 | 1 | 0.76 | 1.01 | 0.61 | 3 |
| acetylcholine-gated monoatomic cation-selective channel activity | 1 | 1 | 1 | 0.76 | 1.01 | 0.61 | 3 |
| ubiquitin-protein transferase activity | 1 | 1 | 1 | 0.76 | 1.01 | 0.61 | 3 |
| serine-type endopeptidase activity | 1 | 1 | 1 | 0.76 | 1.01 | 0.61 | 3 |
| inward rectifier potassium channel activity | 1 | 1 | 1 | 0.76 | 1.01 | 0.61 | 3 |
| molecular adaptor activity | 1 | 1 | 1 | 0.76 | 1.01 | 0.61 | 3 |
| phosphatidylinositol binding | 1 | 1 | 1 | 0.76 | 1.01 | 0.61 | 3 |
| ion channel regulator activity | 1 | 1 | 1 | 0.76 | 1.01 | 0.61 | 3 |
| heparin binding | 1 | 1 | 1 | 0.76 | 1.01 | 0.61 | 3 |
| transcription coregulator activity | 1 | 1 | 1 | 0.76 | 1.01 | 0.61 | 3 |
| hexosyltransferase activity | 1 | 1 | 1 | 0.76 | 1.01 | 0.61 | 3 |
| clathrin light chain binding | 1 | 1 | 1 | 0.76 | 1.01 | 0.61 | 3 |
| dynein light intermediate chain binding | 1 | 1 | 1 | 0.76 | 1.01 | 0.61 | 3 |
| dynein intermediate chain binding | 1 | 1 | 1 | 0.76 | 1.01 | 0.61 | 3 |
| minus-end-directed microtubule motor activity | 1 | 1 | 1 | 0.76 | 1.01 | 0.61 | 3 |
| guanyl-nucleotide exchange factor activity | 1 | 1 | 1 | 0.76 | 1.01 | 0.61 | 3 |
| phospholipid binding | 1 | 1 | 1 | 0.76 | 1.01 | 0.61 | 3 |
| carboxypeptidase activity | 1 | 1 | 1 | 0.76 | 1.01 | 0.61 | 3 |
| syntaxin-1 binding | 1 | 1 | 1 | 0.76 | 1.01 | 0.61 | 3 |
| aminopeptidase activity | 1 | 1 | 1 | 0.76 | 1.01 | 0.61 | 3 |
| metallodipeptidase activity | 1 | 1 | 1 | 0.76 | 1.01 | 0.61 | 3 |
| vitamin transmembrane transporter activity | 1 | 0 | 1 | 0.76 | 0 | 0.61 | 2 |
| vitamin D binding | 1 | 0 | 1 | 0.76 | 0 | 0.61 | 2 |
| G-quadruplex RNA binding | 1 | 0 | 1 | 0.76 | 0 | 0.61 | 2 |
| DNA helicase activity | 1 | 0 | 1 | 0.76 | 0 | 0.61 | 2 |
| transcription cis-regulatory region binding | 1 | 0 | 1 | 0.76 | 0 | 0.61 | 2 |
| structural constituent of postsynaptic intermediate filament cytoskeleton | 1 | 0 | 1 | 0.76 | 0 | 0.61 | 2 |
| C-C chemokine binding | 0 | 1 | 1 | 0 | 1.01 | 0.61 | 2 |
| protein homodimerization activity | 1 | 0 | 1 | 0.76 | 0 | 0.61 | 2 |
| ankyrin binding | 1 | 0 | 1 | 0.76 | 0 | 0.61 | 2 |
| RNA polymerase II transcription regulatory region sequence-specific DNA binding | 0 | 1 | 1 | 0 | 1.01 | 0.61 | 2 |
| thioredoxin peroxidase activity | 0 | 0 | 2 | 0 | 0 | 1.22 | 2 |
| malate synthase activity | 1 | 0 | 1 | 0.76 | 0 | 0.61 | 2 |
| U1 snRNA binding | 1 | 0 | 1 | 0.76 | 0 | 0.61 | 2 |
| U2 snRNA binding | 1 | 0 | 1 | 0.76 | 0 | 0.61 | 2 |
| microtubule binding | 1 | 0 | 1 | 0.76 | 0 | 0.61 | 2 |
| proton-transporting ATP synthase activity, rotational mechanism | 0 | 0 | 2 | 0 | 0 | 1.22 | 2 |
| mRNA binding | 1 | 0 | 1 | 0.76 | 0 | 0.61 | 2 |
| C-C chemokine receptor activity | 0 | 1 | 1 | 0 | 1.01 | 0.61 | 2 |
| structural constituent of muscle | 1 | 0 | 1 | 0.76 | 0 | 0.61 | 2 |
| kinesin binding | 0 | 1 | 0 | 0 | 1.01 | 0 | 1 |
| heme binding | 0 | 0 | 1 | 0 | 0 | 0.61 | 1 |
| catalase activity | 0 | 0 | 1 | 0 | 0 | 0.61 | 1 |
| macrophage colony-stimulating factor receptor activity | 0 | 0 | 1 | 0 | 0 | 0.61 | 1 |
| growth factor binding | 0 | 0 | 1 | 0 | 0 | 0.61 | 1 |
| phosphatidylinositol-3-phosphate binding | 0 | 0 | 1 | 0 | 0 | 0.61 | 1 |
| inositol 1,4,5 trisphosphate binding | 1 | 0 | 0 | 0.76 | 0 | 0 | 1 |
| transferase activity | 1 | 0 | 0 | 0.76 | 0 | 0 | 1 |
| store-operated calcium channel activity | 1 | 0 | 0 | 0.76 | 0 | 0 | 1 |
| phosphatidylserine binding | 0 | 0 | 1 | 0 | 0 | 0.61 | 1 |
| calcium-dependent phospholipid binding | 0 | 0 | 1 | 0 | 0 | 0.61 | 1 |
| lipid transporter activity | 1 | 0 | 0 | 0.76 | 0 | 0 | 1 |
| carbohydrate binding | 0 | 0 | 1 | 0 | 0 | 0.61 | 1 |
| hyaluronic acid binding | 0 | 0 | 1 | 0 | 0 | 0.61 | 1 |
| ATP hydrolysis activity | 1 | 0 | 0 | 0.76 | 0 | 0 | 1 |
| dehydroascorbic acid transmembrane transporter activity | 1 | 0 | 0 | 0.76 | 0 | 0 | 1 |
| diacylglycerol-dependent serine/threonine kinase activity | 1 | 0 | 0 | 0.76 | 0 | 0 | 1 |
| activity | 1 | 0 | 0 | 0.76 | 0 | 0 | 1 |
| ADP binding | 0 | 0 | 1 | 0 | 0 | 0.61 | 1 |
| voltage-gated sodium channel activity | 1 | 0 | 0 | 0.76 | 0 | 0 | 1 |
| porin activity | 0 | 0 | 1 | 0 | 0 | 0.61 | 1 |
| guanine deaminase activity | 1 | 0 | 0 | 0.76 | 0 | 0 | 1 |
| argininosuccinate lyase activity | 0 | 0 | 1 | 0 | 0 | 0.61 | 1 |
| rRNA binding | 0 | 0 | 1 | 0 | 0 | 0.61 | 1 |
| mannosyltransferase activity | 0 | 0 | 1 | 0 | 0 | 0.61 | 1 |

To assess whether EV-based detection reproduces across independent water bodies, EV-DNA and EV-RNA fractions were collected and analysed at Yundang Lake, a separate urban water body, using the same EV isolation and MiFish 12S metabarcoding workflow applied at Xinglinwan Reservoir. As no independent reference species inventory was available for this site, results are reported as detections per EV fraction rather than against an expected-species benchmark, and only the EV fractions are considered. EV-RNA provided broader community coverage than EV-DNA, detecting 11 fish taxa compared with 5 for EV-DNA, and recovering all taxa detected by EV-DNA plus six additional species (including the *cyprinids Cyprinus carpio, Ctenopharyngodon idella, Hypophthalmichthys nobilis, and Megalobrama amblycephala*). Five taxa were detected by both EV fractions. Several of the detected taxa coincided with EV detections at Xinglinwan Reservoir (*Oreochromis niloticus, Clupanodon thrissa, Coptodon zillii, Sarotherodon galilaeus, Ambassis gymnocephalus*), with *O. niloticus* recovered by both EV fractions at both sites, indicating that EV-associated nucleic acids reproducibly recover fish community signals across independent urban water bodies.

**Table S6. Fish taxa detected by EV fractions at Yundang Lake.** *EV-DNA and EV-RNA fractions were sampled at two sites (Y1, Y2; n = 2 per fraction) and processed identically to the Xinglinwan workflow. Values are total read counts per fraction (— = not detected). No independent reference inventory exists for this site, so taxa are listed as detected rather than scored against a benchmark. "Also at XLW" marks taxa likewise detected by EV fractions at Xinglinwan Reservoir.*

| **Species** | **EV-DNA (reads)** | **EV-RNA (reads)** |
| --- | --- | --- |
| *Oreochromis niloticus* | 86 | 203 |
| *Coptodon zillii* | 12 | 151 |
| *Sarotherodon galilaeus* | 11 | 57 |
| *Clupanodon thrissa* | 21 | 27 |
| *Ambassis gymnocephalus* | 8 | 19 |
| *Hypophthalmichthys nobilis* | — | 228 |
| *Cyprinus carpio* | — | 115 |
| *Ctenopharyngodon idella* | — | 81 |
| *Megalobrama amblycephala* | — | 32 |
| *Elopichthys bambusa* | — | 31 |
| *Acentrogobius viridipunctatus* | — | 15 |
